# Using wearable EEG to examine age trends in sleep macro- and micro-architecture across adolescence

**DOI:** 10.1101/2025.10.10.681690

**Authors:** Sanna Lokhandwala, Rebecca Cooper, Rebecca Hayes, Soumya Sathe, Isabelle Elder, Mary Corcoran, Beatriz Horta, Maya Fray-Witzer, Lauren Keller, Simey Chan, Peter Franzen, Daniel Buysse, Brant P. Hasler, Jessica C. Levenson, Meredith L. Wallace, Duncan B Clark, Ronette G. Blake, Adriane Soehner, Maria Jalbrzikowski

**Author notes:** Corresponding Author: Maria Jalbrzikowski, Ph.D., Department of Psychiatry, Boston Children’s Hospital, 1 Brookline Place, Suite 335, Brookline, MA 02445.

## Abstract

Adolescence is a period of distinct maturational changes in sleep physiology. Age-related trends in sleep physiology have been captured using laboratory-based polysomnography, a method limited by logistical burden and high cost. We tested the ability of the Dreem3 sleep EEG headband to replicate established age effects in sleep physiology from late childhood through early adulthood. Typically developing youth (N=100, 9-26 years) completed 3-4 consecutive nights of at-home sleep recording. We estimated age-related trends across eight macro-architecture and 15 micro-architecture variables with known age effects and conducted exploratory analyses of 24 additional variables. The Dreem3 replicated some established age trends, including increases in non-rapid eye movement (NREM) stage 2%, and decreases in N3%, time in bed, NREM delta and theta power with increasing age. Although not all expected age-related associations reached statistical significance, like total sleep time and sleep efficiency, effect estimates were largely in the expected direction. These findings highlight the promise of wearable EEG devices for characterizing developmental sleep physiology while underscoring the need for further validation across sleep measures and age groups.

## INTRODUCTION

The transition from late childhood to adulthood is a unique developmental period marked by brain maturation and sleep changes. Sleep physiology features, which are sensitive markers of brain development, undergo prominent transformations in the second decade of life^1,2^. Sleep physiology plays an active role in sculpting the maturation of the brain^3^, predicts long-term brain, behavior, and health trajectories^4–6^, and is modifiable with non-invasive biobehavioral intervention^7–10^. Delineating the physiological sleep changes that occur over adolescence in a scalable way is critical for understanding developmental variability and how aberrant individual-level health trajectories manifest.

Sleep macro- and micro-architecture undergo dramatic changes throughout adolescence^1,11,12^. Sleep macro-architecture is based on standard sleep staging via polysomnography (PSG) and refers to the overall structure and organization of sleep across the entire sleep period. Total sleep time (TST), percent of time spent in non-rapid eye movement (NREM) stage 3 sleep (N3%), and rapid-eye movement (REM) latency decrease with increasing age^13–17^. In contrast, there is a concomitant rise in percent of time spent in NREM stage 2 sleep (N2%) and wake after sleep onset (WASO)^14,18,19^. Sleep micro-architecture captures specific electrophysiological features within different sleep stages^18^. Cross-sectional studies find significant age effects in slow wave sleep characteristics, showing age-associated decreases in NREM delta power, and slow wave amplitude and slope^11,12,20,21^. Longitudinal studies have further confirmed this decline across adolescence^15,22^. For instance, one longitudinal study of adolescents 9 to 17 years of age showed that NREM delta power (1-4 Hz) exhibited a steep decline between 11 and 12 years^22^. Other studies find that sigma power (12-15 Hz) and various spindle characteristics increase over adolescence^18,23–26^. For example, Kozhemiako and colleagues (2024) found that absolute sigma power and fast spindle density increased with age^18^. These developmental shifts in sleep EEG patterns during adolescence are linked to brain maturational processes which influence long-term brain, behavior, and health trajectories^4–6^.

Age trends in sleep macro- and micro-architecture have historically been captured using laboratory-based PSG. Multiple challenges associated with PSG make it prohibitive to measure age effects for large-scale use and incorporate sleep EEG measurement into community settings. Standard PSG is costly, requires highly trained staff, and is time-consuming to set up, conduct, and score, making it impractical for multi-night home-based recordings on a large scale. Generalizability of lab-based PSG is also a concern, as in-lab PSG has known ‘first night effects’ due to sleeping in a new environment^27^. First night effects are characterized by shorter and more fragmented sleep^27^, potentially preventing researchers from reliably capturing an individuals’ habitual sleep patterns, raising questions about the ecological validity of PSG.

Among adolescents in particular, multiple nights of sleep recording may be needed to obtain stable and valid sleep EEG estimates^28^. Meta-analytic and longitudinal data point to weekday-weekend differences in sleep EEG characteristics in adolescents^15,19^. Further, sleep loss, irregularity, fragmentation, or timing shifts, common occurrences in adolescents^29^, can introduce variability in sleep stage duration and sleep micro-architecture^30^, reducing the reliability of sleep endpoints drawn from a single-night recording. Indeed, measures of within-subject variability from at-home wearable sleep devices may more accurately capture adolescent sleep than laboratory PSG^31^. In a sample of adults, Arnal and colleagues (2020) found that an at-home, wireless sleep monitoring device, the Dreem headband, acquired EEG signals that strongly correlated with EEG signals from PSG^32^. Furthermore, the Dreem automatic sleep staging algorithm performed at a similar level of accuracy to expert scorers^32^. Similarly, in a recent study of adolescents and young adults, the Dreem device provided sleep estimates that are moderately stable between nights, and these estimates are similar between youth and adults^31^. These results highlight that adopting a more portable, inexpensive, and scalable approach to sleep EEG results in similar outcomes as PSG for the sleep field^33^.

It is critical to translate sleep-based risk assessment from research settings to community settings to have the greatest impact^34,35^. By incorporating sleep assessment into community settings, we could identify those at highest risk (e.g., for psychiatric illness) earlier, reach individuals traditionally underserved by academic medical centers, and provide a more accurate picture of age trends in the general population^36,37^. Using accessible, in-home sleep measures not only satisfies service users’ preference for community settings and easily implemented assessments^36^ but also removes logistical barriers that prevent individuals (often under-represented or minoritized) from participating in research or seeking clinical care, such as lack of transportation, difficulty finding childcare, or lengthy time commitments.

To this end, the current study aims to address these theoretical and technical gaps by attempting to replicate known age-related trends in sleep macro- and micro-architecture in young people using the Dreem3 headband. We considered sleep measures as having a “replicated” age effect if at least two independent studies reported consistent relationships in the same direction (See Tables S1 & S2). Specifically, we examined eight sleep macro-architecture features: TST, WASO, N2%, N3%, REM%, REM latency, sleep efficiency (SE), and time in bed (TIB)^11,13–15,17,18,38,39^. We examined fifteen sleep micro-architecture features, including NREM and REM spectral power (absolute delta, sigma, theta, and beta power) and oscillatory characteristics (slow spindle amplitude, density, duration; fast spindle amplitude, density, and duration; and slow oscillation-spindle coupling magnitude for slow and fast spindles)^3,11,13,20–23,38,40–45^. We explored 24 additional sleep macro- and micro-architecture features with less established age effects (i.e., no study has examined age associated trends in these features or we did not find at least two papers showing consistent age effects). Finally, we assessed age-associated trends in night-to-night variability in macro- and micro-architectural features within individuals.

## RESULTS

### Sample description

The final sample comprised 100 participants (53% female) with 309 nights of data (mean number of nights: 3.09, standard deviation of number of nights: 1.04, and range of number of nights 1-9). See Table 1 for sample demographics and Table S3 for descriptive statistics of the sleep variables examined.

**Table 1.**
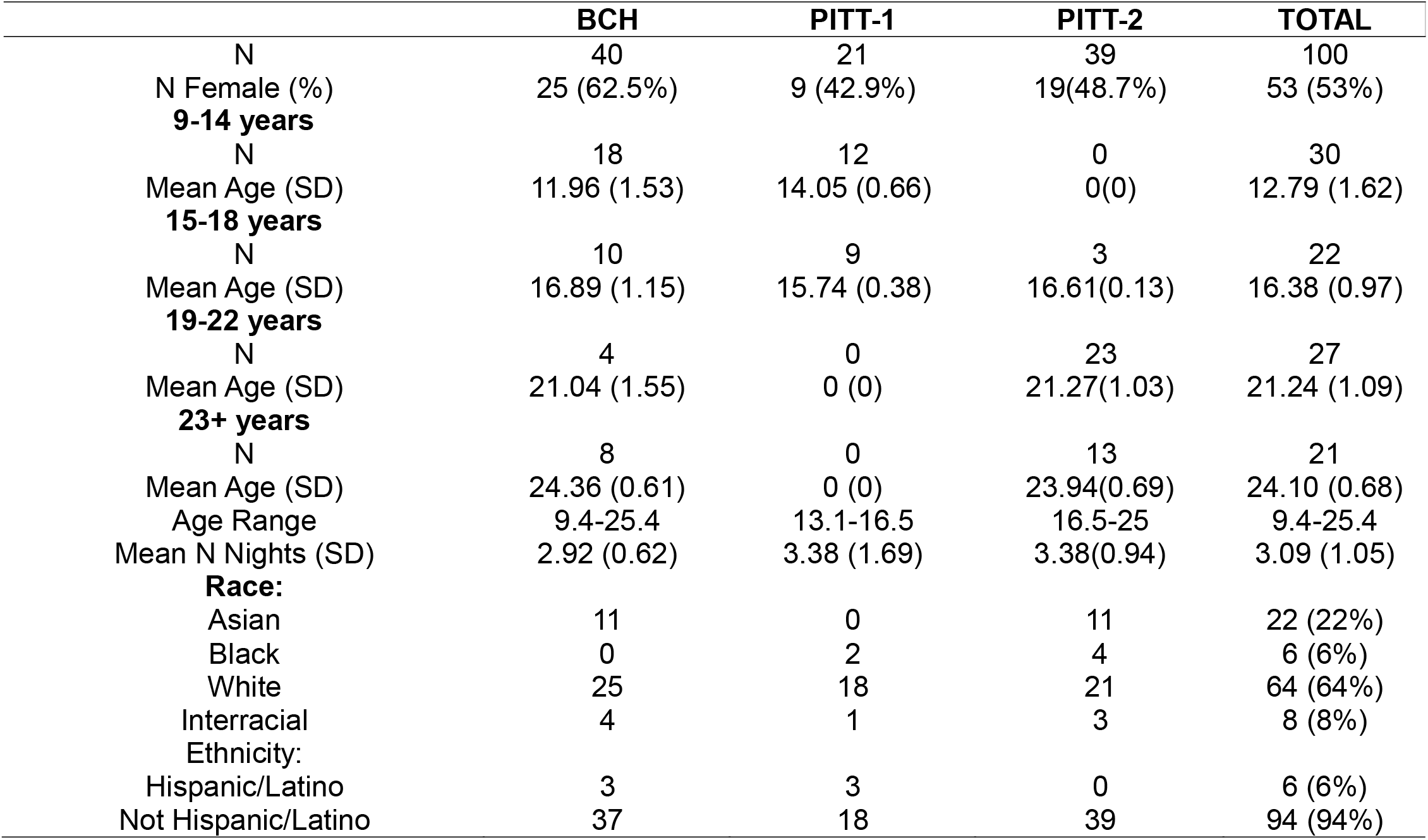
Sample demographics.

|  | **BCH** | **PITT-1** | **PITT-2** | **TOTAL** |
|---|---|---|---|---|
| N | 40 | 21 | 39 | 100 |
| N Female (%) | 25 (62.5%) | 9 (42.9%) | 19(48.7%) | 53 (53%) |
| **9-14 years** |  |  |  |  |
| N | 18 | 12 | 0 | 30 |
| Mean Age (SD) | 11.96 (1.53) | 14.05 (0.66) | 0(0) | 12.79 (1.62) |
| **15-18 years** |  |  |  |  |
| N | 10 | 9 | 3 | 22 |
| Mean Age (SD) | 16.89 (1.15) | 15.74 (0.38) | 16.61(0.13) | 16.38 (0.97) |
| **19-22 years** |  |  |  |  |
| N | 4 | 0 | 23 | 27 |
| Mean Age (SD) | 21.04 (1.55) | 0 (0) | 21.27(1.03) | 21.24 (1.09) |
| **23+ years** |  |  |  |  |
| N | 8 | 0 | 13 | 21 |
| Mean Age (SD) | 24.36 (0.61) | 0 (0) | 23.94(0.69) | 24.10 (0.68) |
| Age Range | 9.4-25.4 | 13.1-16.5 | 16.5-25 | 9.4-25.4 |
| Mean N Nights (SD) | 2.92 (0.62) | 3.38 (1.69) | 3.38(0.94) | 3.09 (1.05) |
| **Race:** |  |  |  |  |
| Asian | 11 | 0 | 11 | 22 (22%) |
| Black | 0 | 2 | 4 | 6 (6%) |
| White | 25 | 18 | 21 | 64 (64%) |
| Interracial | 4 | 1 | 3 | 8 (8%) |
| Ethnicity: |  |  |  |  |
| Hispanic/Latino | 3 | 3 | 0 | 6 (6%) |
| Not Hispanic/Latino | 37 | 18 | 39 | 94 (94%) |

### Replication of known age effects with at-home EEG device in adolescents - sleep macro-architecture

We observed significant age-associated increases in N2% (β = 0.28, *p*_FDR_ = 7.91e-04), and age-associated decreases in N3% (β = -0.42, *p*_FDR_ = 5.82e-07), REM latency (β = -0.25, *p*_FDR_ = 3.10e-04) and time in bed (β = -0.25, *p*_FDR_ = 7.91e-04; Figure 1; Table 2). However, we did not find age-related associations with TST (Figure 1), WASO, REM%, and sleep efficiency (all *p*_FDR_ ≥ 0.11), although effect size estimates were generally in the expected direction (Table 2). There was no main effect of sex for any of the eight macro-architecture variables following false discovery rate (FDR) correction (all *p*_FDR_>0.114, Figure S1). Post hoc sensitivity analyses excluding all in-lab recording nights yielded largely unchanged results. However, within these sensitivity analyses, we also observed an additional statistically significant effect of age on percent of time spent in REM sleep, with an increase in percentage time spent in REM with increasing age (β = 0.16, *p*_FDR_ = 2.14e-02; Table S4). Further, results were consistent when accounting for the effect of weekday/weekend night status or summer/school season (Tables S6 & S7).

**Figure 1.**
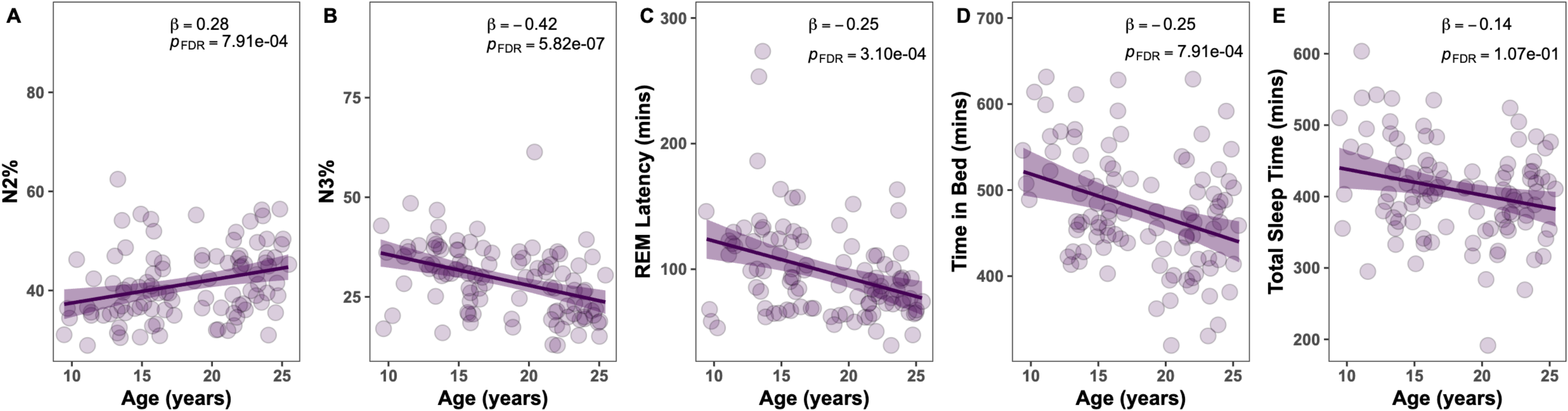
Associations between sleep macro-architecture and age. (A) non-rapid eye movement (NREM) stage 2 (N2%) sleep, (B) NREM stage 3 (N3%) sleep, (C) rapid-eye movement (REM) latency, (D) time in bed, (E) total sleep time. For visualization purposes, we averaged data points across channel and night for each individual. Line represents regression line for model; shading represents 95% confidence intervals.

**Table 2.**
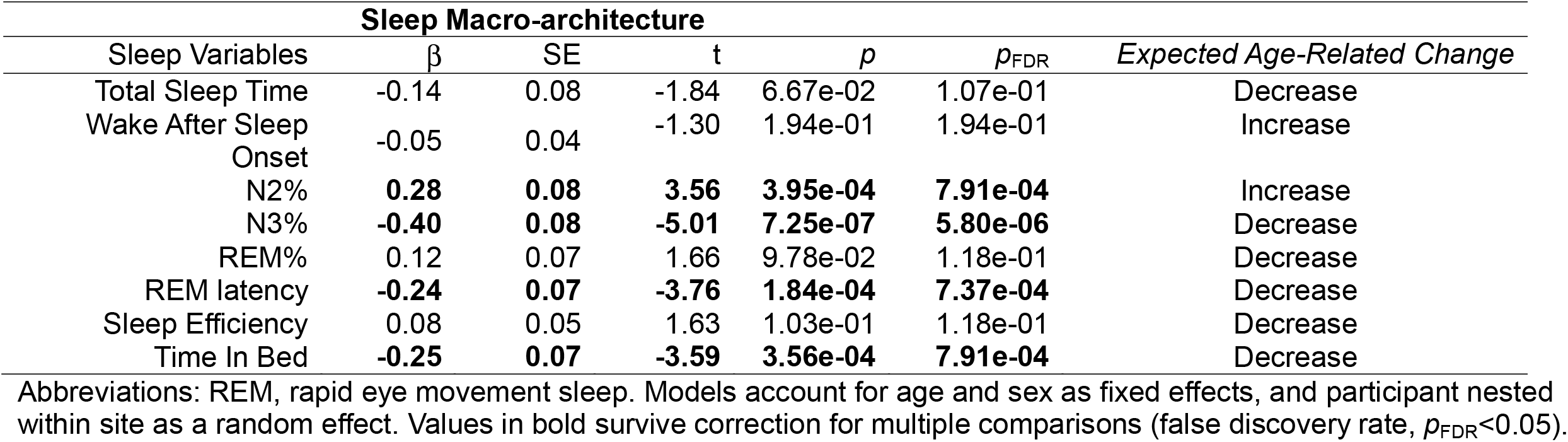
Linear mixed-effects model for the effect of age on sleep macro-architecture across adolescence.

| **Sleep Macro-architecture** |  |  |  |  |  |
|---|---|---|---|---|---|
| Sleep Variables | β | SE | t | *p* | *p*<sub>FDR</sub> | *Expected Age-Related Change* |
| Total Sleep Time | -0.14 | 0.08 | -1.84 | 6.67e-02 | 1.07e-01 | Decrease |
| Wake After Sleep Onset | -0.05 | 0.04 | -1.30 | 1.94e-01 | 1.94e-01 | Increase |
| N2% | **0.28** | **0.08** | **3.56** | **3.95e-04** | **7.91e-04** | Increase |
| N3% | **-0.40** | **0.08** | **-5.01** | **7.25e-07** | **5.80e-06** | Decrease |
| REM% | 0.12 | 0.07 | 1.66 | 9.78e-02 | 1.18e-01 | Decrease |
| REM latency | **-0.24** | **0.07** | **-3.76** | **1.84e-04** | **7.37e-04** | Decrease |
| Sleep Efficiency | 0.08 | 0.05 | 1.63 | 1.03e-01 | 1.18e-01 | Decrease |
| Time In Bed | **-0.25** | **0.07** | **-3.59** | **3.56e-04** | **7.91e-04** | Decrease |

Abbreviations: REM, rapid eye movement sleep. Models account for age and sex as fixed effects, and participant nested within site as a random effect. Values in bold survive correction for multiple comparisons (false discovery rate, *p*<sub>FDR</sub><0.05).

### Replication of known age effects with at-home EEG device in adolescents - sleep micro-architecture

Our models revealed NREM delta power (β = -0.49, *p*_FDR_ = 1.00e-11) and NREM theta power (β = -0.10, *p*_FDR_ = 9.15e-03) decreased with increasing age. Slow spindle duration (β = -0.41, *p*_FDR_ = 4.50e-12), slow spindle density (β = -0.43, *p*_FDR_ = 3.83e-07), and fast spindle duration (β = -0.17, *p*_FDR_ = 6.92e-05) also decreased with increasing age. Lastly, the magnitude of slow oscillation-spindle coupling for slow (β = 0.15, *p*_FDR_ = 2.01e-03) and fast (β = 0.10, *p*_FDR_ = 1.01e-02) spindles increased with increasing age (Figure 2, Table 3). However, we did not find age associated effects in absolute NREM sigma, beta, REM delta, sigma, theta power (p_FDR_ ≥ 5.50e-01), slow spindle amplitude, fast spindle amplitude, and fast spindle density (p_FDR_ ≥ 4.57e-01), although most effect size estimates were in the expected direction (Table 3, Figure S4).

**Figure 2.**
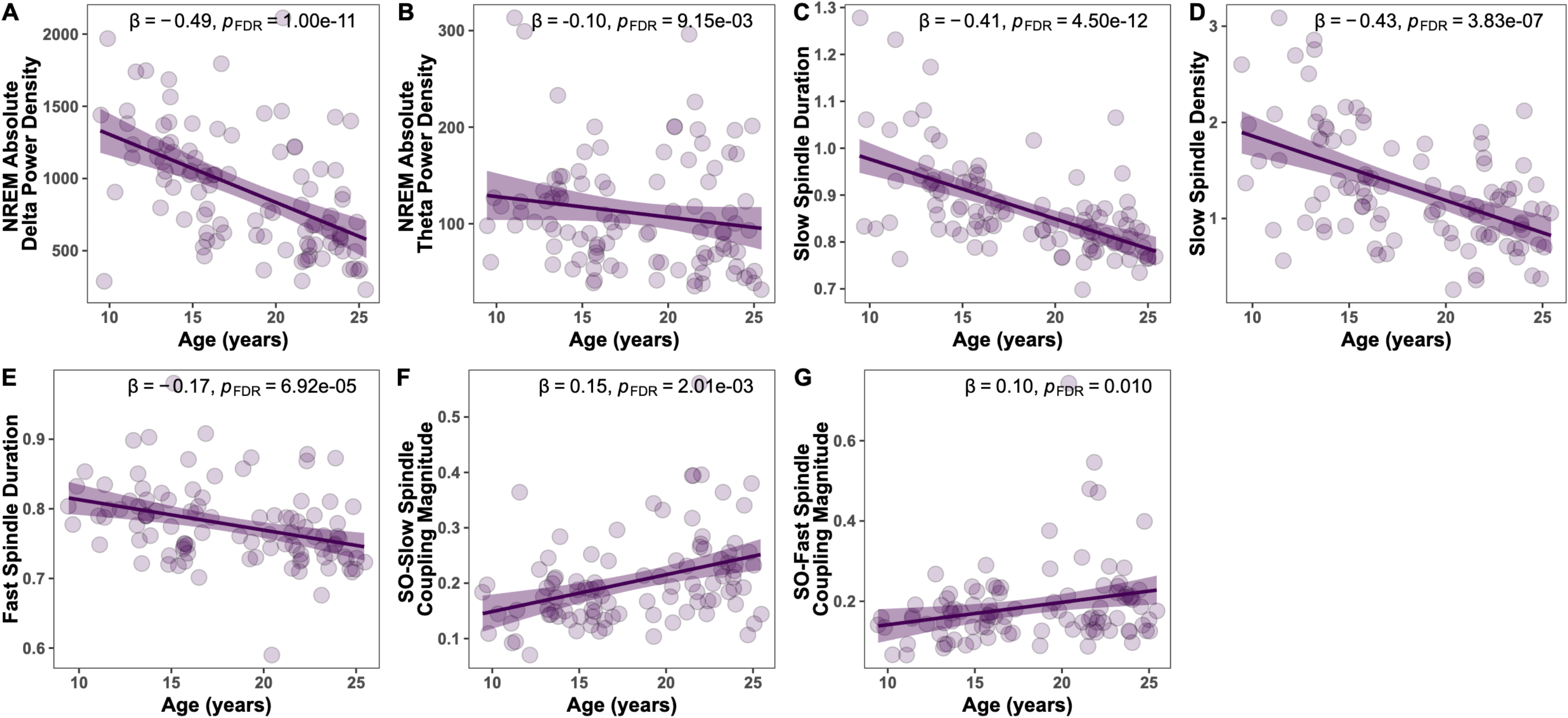
Associations between sleep micro-architecture and age. (A) non-rapid eye movement (NREM) absolute delta power density, (B) NREM absolute theta power density, (C) slow spindle duration, (D) slow spindle density, (E) fast spindle duration, (F) slow oscillation (SO)-slow spindle coupling magnitude, (G) SO-fast spindle coupling magnitude. For visualization purposes, we averaged data points across channel and night for each individual. Line represents regression line for model; shading represents 95% confidence intervals.

**Table 3.**
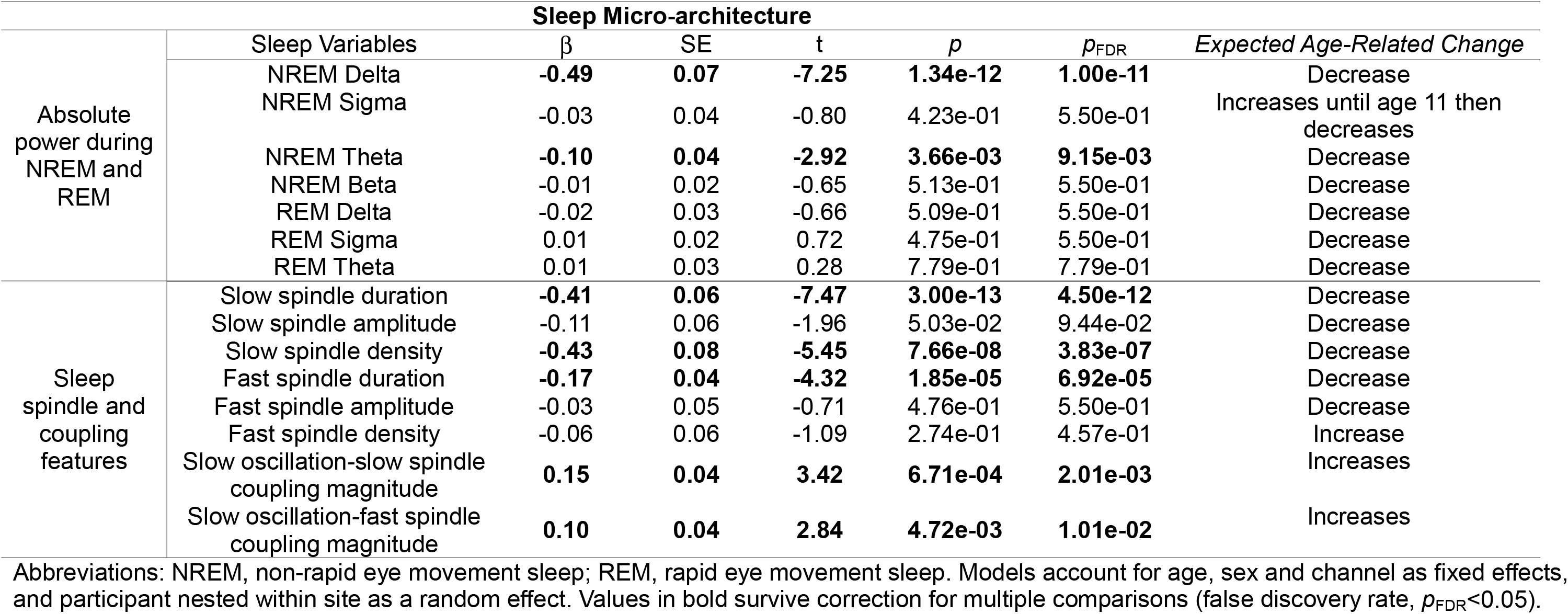
Linear mixed-effects model for the effect of age on sleep micro-architecture across adolescence.

| Sleep Micro-architecture |  |  |  |  |  |  |  |
|---|---|---|---|---|---|---|---|
|  | Sleep Variables | β | SE | t | *p* | *p*<sub>FDR</sub> | *Expected Age-Related Change* |
| Absolute power during NREM and REM | NREM Delta | **-0.49** | **0.07** | **-7.25** | **1.34e-12** | **1.00e-11** | Decrease |
|  | NREM Sigma | -0.03 | 0.04 | -0.80 | 4.23e-01 | 5.50e-01 | Increases until age 11 then decreases |
|  | NREM Theta | **-0.10** | **0.04** | **-2.92** | **3.66e-03** | **9.15e-03** | Decrease |
|  | NREM Beta | -0.01 | 0.02 | -0.65 | 5.13e-01 | 5.50e-01 | Decrease |
|  | REM Delta | -0.02 | 0.03 | -0.66 | 5.09e-01 | 5.50e-01 | Decrease |
|  | REM Sigma | 0.01 | 0.02 | 0.72 | 4.75e-01 | 5.50e-01 | Decrease |
|  | REM Theta | 0.01 | 0.03 | 0.28 | 7.79e-01 | 7.79e-01 | Decrease |
| Sleep spindle and coupling features | Slow spindle duration | **-0.41** | **0.06** | **-7.47** | **3.00e-13** | **4.50e-12** | Decrease |
|  | Slow spindle amplitude | -0.11 | 0.06 | -1.96 | 5.03e-02 | 9.44e-02 | Decrease |
|  | Slow spindle density | **-0.43** | **0.08** | **-5.45** | **7.66e-08** | **3.83e-07** | Decrease |
|  | Fast spindle duration | **-0.17** | **0.04** | **-4.32** | **1.85e-05** | **6.92e-05** | Decrease |
|  | Fast spindle amplitude | -0.03 | 0.05 | -0.71 | 4.76e-01 | 5.50e-01 | Decrease |
|  | Fast spindle density | -0.06 | 0.06 | -1.09 | 2.74e-01 | 4.57e-01 | Increase |
|  | Slow oscillation-slow spindle coupling magnitude | **0.15** | **0.04** | **3.42** | **6.71e-04** | **2.01e-03** | Increases |
|  | Slow oscillation-fast spindle coupling magnitude | **0.10** | **0.04** | **2.84** | **4.72e-03** | **1.01e-02** | Increases |

Abbreviations: NREM, non-rapid eye movement sleep; REM, rapid eye movement sleep. Models account for age, sex and channel as fixed effects, and participant nested within site as a random effect. Values in bold survive correction for multiple comparisons (false discovery rate, *p*<sub>FDR</sub><0.05).

There was no main effect of sex for any of the fifteen sleep microarchitecture variables following FDR correction (all *p*_FDR_>0.866; Figure S2). Post hoc sensitivity analyses excluding all in-lab recording nights returned similar results to our original analyses (Table S5). Further, results were consistent when examining the effect of weekday/weekend night or summer/school season (Table S8 & S9).

### Exploratory analyses

We also explored age effects in 24 additional macro- and micro-architecture variables. After multiple comparison correction, we found statistically significant age-associated trends for twelve measures. We observed age-associated increases in slow oscillation duration (β = 0.21, *p*_FDR_ = 0.018), relative NREM sigma (β = 0.23, *p*_FDR_ = 1.03e-04), relative NREM beta (β = 0.12, *p*_FDR_ = 9.10e-07), relative REM sigma (β = 0.15, *p*_FDR_ = .028), relative REM theta (β = 0.13, *p*_FDR_ = .030), and relative REM alpha power (β = 0.21, *p*_FDR_= 1.22e-03). We observed age-associated decreases in total NREM cycle duration (β = -0.21, *p*_FDR_ = 0.024), median slow oscillation peak to peak amplitude (β = -0.49, *p*_FDR_= 8.50e-12), relative NREM delta power (β = -0.27, *p*_FDR_ = 8.63e-03), relative REM delta power (β = -0.15, *p*_FDR_ = .031), and integrated slow (β = -0.35, *p*_FDR_= 2.71e-07) and fast spindle activity (β = -0.12, *p*_FDR_ = 0.032) with increasing age. The full list of results is reported in Table 4.

**Table 4.** Exploratory: Linear mixed-effects model for the effect of age on sleep macro- and micro-architecture with lesser-known age effects across adolescence.

| Sleep Macro- and Micro-architecture |  |  |  |  |  |  |
|  | Sleep Variables | β | SE | t | *p* | *p*<sub>FDR</sub> |
|---|---|---|---|---|---|---|
| Sleep macro-architecture features | Transition index from NREM to REM | -0.03 | 0.08 | -0.38 | 7.06e-01 | 7.06e-01 |
|  | Total sleep cycles | -0.06 | 0.06 | -0.89 | 3.73e-01 | 6.01e-01 |
|  | Total cycle duration | -0.13 | 0.08 | -1.65 | 9.99e-02 | 2.50e-01 |
|  | Total NREM cycle duration | **-0.21** | **0.08** | **-2.83** | **4.86e-03** | **2.43e-02** |
|  | Total REM cycle duration | 0.06 | 0.08 | 0.70 | 4.81e-01 | 6.01e-01 |
| Slow oscillation features | Slow oscillation duration | **0.21** | **0.08** | **2.73** | **6.48e-03** | **1.76e-02** |
|  | Median slow oscillation peak to peak amplitude | **-0.49** | **0.07** | **-7.41** | **4.47e-13** | **8.50e-12** |
| Relative power during NREM and REM | NREM Delta | **-0.27** | **0.09** | **-3.01** | **2.73e-03** | **8.63e-03** |
|  | NREM Sigma | **0.23** | **0.05** | **4.28** | **2.17e-05** | **1.03e-04** |
|  | NREM Theta | 0.11 | 0.05 | 1.94 | 5.31e-02 | 8.41e-02 |
|  | NREM Beta | **0.12** | **0.02** | **5.33** | **1.44e-07** | **9.10e-070** |
|  | REM Delta | **-0.15** | **0.06** | **-2.41** | **1.63e-02** | **3.10e-02** |
|  | REM Sigma | **0.15** | **0.06** | **2.53** | **1.16e-02** | **2.75e-02** |
|  | REM Theta | **0.13** | **0.05** | **2.46** | **1.42e-02** | **2.99e-02** |
|  | REM Alpha | **0.21** | **0.06** | **3.62** | **3.22e-04** | **1.22e-03** |
|  | REM Beta | 0.05 | 0.03 | 1.51 | 1.31e-01 | 1.66e-01 |
| Absolute power during REM | REM Alpha | 0.03 | 0.02 | 1.38 | 1.67e-01 | 1.87e-01 |
|  | REM Beta | 0.00 | 0.01 | 0.13 | 8.97e-01 | 8.97e-01 |
| Sleep spindle features | Integrated slow spindle activity | **-0.35** | **0.06** | **-5.63** | **2.85e-08** | **2.71e-07** |
|  | Integrated fast spindle activity | **-0.12** | **0.05** | **-2.36** | **1.85e-02** | **3.20e-02** |
|  | Slow spindle chirp | 0.14 | 0.09 | 1.68 | 9.27e-02 | 1.35e-01 |
|  | Fast spindle chirp | -0.09 | 0.06 | -1.47 | 1.42e-01 | 1.68e-01 |
|  | Slow spindle frequency | -0.05 | 0.09 | -0.56 | 5.74e-01 | 6.06e-01 |
|  | Fast spindle frequency | 0.14 | 0.09 | 1.64 | 1.02e-01 | 1.39e-01 |

Abbreviations: PSD, absolute spectral power density; RELPSD, relative spectral power density. Models account for age and sex as fixed effects, and participant nested within site as a random effect. Microarchitecture variable models also included channel as fixed effect.

#### Sex differences

We observed significant sex effects for one exploratory macro-architecture and three exploratory micro-architecture variables. In comparison to males, females had a significantly greater total number of sleep cycles (β = -0.37, *p*_FDR_ = 0.016). However, males had higher relative REM sigma power (β = 0.36, *p*_FDR_ = 0.015), relative REM theta power (β = 0.34, *p*_FDR_ = 0.015), and relative REM beta power (β = 0.17, *p*_FDR_ = 0.025) in comparison to females. We did not observe any statistically significant age-by-sex interactions (see Table S10 and Figure S3).

#### Intra-individual variability

We observed increasing age was associated with increased night-to-night variability in REM% (β = 0.21, *p*_FDR_ = 0.011), slow oscillation-slow spindle coupling magnitude (β = 0.25, *p*_FDR_ = 0.032), relative NREM theta power (β = 0.17, *p*_FDR_ = 0.039), relative NREM beta power (β = 0.03, *p*_FDR_ = 0.039), and slow spindle chirp (β = 0.25, *p*_FDR_ =0.039; See Table S11 and Figure S5. Original intra-individual variability analyses included both at-home and in-lab recordings. In post-hoc sensitivity analyses restricted to at-home recordings, the original age-related associations were no longer statistically significant, although the directionality of the effects remained consistent and effect sizes were similar (Table S12). Indeed, βs from the original variability analyses were strongly correlated with the βs from the at-home-only variability analyses (r=.87, *p*=1.80e-15). Additionally, within-subject paired-samples t-tests revealed no significant differences between home and in-lab recordings across the examined macro- and micro-architecture variables (*p*_FDR_s>0.427; Table S13). Within the at-home-only intra-individual variability analyses, we observed that increasing age was associated with decreased night-to-night variability in REM latency (β = 0.-40, *p*_FDR_ = 4.65e-04) and decreased night-to-night variability in absolute NREM delta power (β = 0.-29, *p*_FDR_ = 1.85e-02; Table S12).

## DISCUSSION

We assessed age-associated trends in sleep behavior across adolescence using an accessible, at-home sleep EEG device. Using the Dreem3 headband, we captured macro- and micro-architecture age effects previously characterized using lab-based PSG. We were able to replicate known age effects in macro-architecture variables N2%, N3%, REM latency and time in bed. Further, in line with previous PSG studies examining age trends in micro-architecture, we found age-related associations with delta and theta power and spindle characteristics (e.g., slow and fast spindle duration). We explored age-related effects in macro- and micro-level variables with limited prior evidence of age associations (e.g., slow and fast spindle frequency), revealing several additional age-related relationships. Finally, we identified age-associated increases in night- to-night variability in several macro- and micro-architectural features (e.g., REM%, NREM beta and theta power). Taken together, these results suggest many of the known age effects observed in lab-based PSG in youth can be captured through at-home wearable sleep EEG recordings, and specifically, with the Dreem3 headband. Furthermore, wearable devices enable researchers to explore lesser-known age effects associated with other sleep physiology variables at scale. These results demonstrate the potential for using at-home sleep EEG wearables in young people and provide a foundation for linking sleep physiology measures to behavioral and mental health trajectories.

Of the 23 macro- and micro-architecture sleep variables with replicated age associations (i.e., at least two independent PSG studies reported consistent relationships in the same direction), we found significant age effects in approximately half of these features. Similar to PSG studies^11,13,14,18,38,46^, we found N2% increased with age while N3%, REM latency and time in bed decreased with age. For micro-architecture, we found that slow oscillation-slow and -fast spindle coupling magnitude increased with age, while absolute NREM delta and theta power, slow and fast spindle duration, and slow spindle density decreased with age. Replicating age effects observed in PSG studies provides additional evidence that the Dreem3 headband is capturing physiologically meaningful EEG features. Thus, at-home wearable sleep EEG headbands offer a valuable alternative to lab-based PSG in understanding developmental effects on sleep physiology. Additionally, wearables allow researchers and clinicians to capture longitudinal and ecologically valid data, as such devices allow for multi-night, at-home recordings, which PSG rarely affords due to cost and logistical constraints. This makes wearables ideal for studying not only group-level comparisons but within-person effects as well.

We did not replicate several age effects reported in PSG studies. For example, we did not replicate age effects of TST, WASO, sleep efficiency, NREM beta and sigma power, and several REM spectral and spindle features. The absence of a statistically significant age-related association with TST is particularly notable given prior studies reporting age-related declines in TST^14,18^, although we did observe a statistically significant age-related decline in TIB. However, the estimated association between age and TST is in the expected negative direction and comparable in magnitude to previous work^18^, highlighting the need for large samples when measuring age effects with at-home EEG wearables. We considered whether laboratory recording protocols might constrain sleep duration. However, sensitivity analyses excluding participants with in-lab recordings yielded largely unchanged results, suggesting that the inclusion of laboratory recordings did not account for this finding. Furthermore, within-subject comparisons of laboratory and at-home recordings did not suggest that recording context systematically accounted for the observed patterns across established macro- and micro-architecture measures.

Although the present study largely captured naturalistic sleep patterns using wearable EEG in the home environment, prior developmental studies have reported age-related declines in TST using home-based ambulatory PSG (e.g.,^15^), suggesting differences in recording environment alone are unlikely to explain the discrepancies. Differences in recording devices and study protocols between the current study and prior developmental studies^14,15,18^, sample characteristics, and/or analytical approaches may contribute to these discrepancies. Altogether, these findings warrant caution when interpreting the Dreem3’s ability to capture established developmental changes in sleep duration and other aspects of sleep physiology.

Furthermore, it is important to consider the influence of eye-movement artifacts on frontal EEG recordings when characterizing REM EEG using the Dreem3. Our current version of the processing pipeline does not specifically remove or regress out eye-movements from REM EEG. Although excluding EEG segments containing eye movements may reduce ocular contamination, this approach may also limit analyses to tonic REM sleep and disproportionately exclude periods of phasic REM sleep^47^, thereby limiting the range of REM physiology available for analysis. Our future work using simultaneously acquired Dreem EEG and PSG data will provide an opportunity to refine methods for identifying and regressing out these artifacts, ultimately enhancing our ability to measure REM signals. Further, while our design may better capture real-world sleep (e.g., allowing for self-selected bedtimes and wake times), it can also magnify the influence of certain confounds, such as control of ambient light and screen exposure before bedtime. The discrepancies between findings from at-home vs. in-lab sleep studies may reveal important differences between sleep capacity (i.e., more controlled and standardized conditions in-lab) and sleep expression (i.e., more ecologically valid but less controlled home studies).

In contrast to PSG, Dreem3 has dry (and fewer) EEG electrodes and relies on a proprietary automated algorithm for sleep staging. Dry-EEG electrodes embedded in the Dreem3 headband may introduce variability in electrode contact and impedance. In turn, a noisy signal may not only compromise WASO and sleep efficiency estimations (e.g., masking alpha activity) but influence spectral power as well^48^, which may explain why we only observed age effects in two frequency bands (i.e., NREM delta and theta). The absence of expected age-related associations with some REM and spindle features may reflect limitations of Dreem’s automated staging algorithm. Dreem has shown only moderate agreement between devices and PSG for multi-stage classification and some EEG band power (i.e., sigma power), and an overestimation of REM^49^. Algorithmic misclassifications of REM and sigma power could attenuate detection of age-related effects in REM and spindle features. Discrepancies with prior PSG studies may also derive from fewer electrode sites in the Dreem3 headband. The Dreem3 headband records from frontal and occipital sites while PSG records from multiple derivations, including central and parietal, where spectral activity may be more reliably detected. Specifically, PSG typically detects age-related effects in sleep using central derivations such as C3-M2 and C4-M1^50,51^. For instance, fast spindles are most prominent over central and parietal regions^52^); thus, the device’s frontal-weighted montage may underestimate the fast spindles relative to slow spindles, as slow spindles are observed more frontally. Different referencing schemes also impact EEG micro-architecture. For instance, Dreem3 references the frontal channels to O1 and O2 (i.e., bipolar recording) which could introduce more alpha and attenuate overall signal amplitude. In contrast, traditional PSG typically references channels to the contralateral mastoid (i.e., M1/M2; referential recording) because they are considered relatively ‘electrically neutral.’ More generally, Dreem3 may be more prone to artifact due to limited impedance control and greater susceptibility to the impact of movement, sweat, and hair on the EEG signal, which can distort physiological differences (e.g., more high frequency signal related to electrodes slipping off the head)^53^. Considering these differences, future studies could incorporate measurement error modeling to build a functional relationship between the “gold-standard”, more accurate measure (i.e., PSG) and the lower cost alternative, which is also less precise (i.e., Dreem3 headband) and then use this functional relationship to improve our ability to calculate relationships between the lower cost measure and various outcomes^54,55^.

We also explored age-related associations in several less frequently explored sleep measures. Of 24 measures, we observed age associations with 12 variables. We found age-associated decreases in total NREM cycle duration, median slow oscillation peak to peak amplitude, relative NREM and REM delta power, and integrated slow and fast spindle activity. We also observed age-associated increases in relative NREM sigma and relative REM sigma, theta and alpha power. Interestingly, we found more age associations with relative power than absolute power across most frequency bands. It may be that relative power remains robust across measurement methods because it is internally normalized (e.g., sigma power as a percentage of total power). These results suggest that while certain age-related changes in sleep are well-documented, novel findings in other variables could point to other aspects of NREM and REM sleep that change with age in previously unrecognized yet meaningful ways.

We found that night-to-night variability increased with age across five sleep features: REM%, slow-oscillation (SO)-slow spindle coupling magnitude, relative NREM theta and beta power and slow spindle chirp. Prior work using actigraphy has shown that adolescence is a period of increasing irregularity in sleep macro-architectural features - particularly sleep duration and timing - peaking in late adolescence to early adulthood and declining thereafter^56–59^. For micro-architectural features, prior work in children and adolescents has shown moderate to high stability (intra-class correlation coefficients >0.70) in NREM power spectra across nights^46,60,61^; however, to our knowledge, no studies have directly examined whether this stability varies with age in young people. In adults, coupling between slow wave oscillations and slow frequency spindles appears highly stable within individuals (intra-class correlation coefficients >0.70)^62,63^; however, in female adults, REM% appears more variable (intra-class correlations 0.31-0.39)^64^. An important consideration when interpreting the present night-to-night variability findings is that the main analyses included a combination of at-home and in-lab recordings. When we conducted sensitivity analyses restricting the analyses to at-home recordings only, the original age-related associations were no longer statistically significant. However, the directionality of these effects remained consistent and effect sizes were very similar. Indeed, βs from the original variability analyses were highly correlated with the βs from the at-home-only models (r=.87, *p*=1.80e-15). Additionally, within-subject comparisons revealed no significant differences between home and in-lab recordings across sleep variables, suggesting that sleep measures from in-lab recordings were not quantitatively different from the sleep measures obtained from at-home recordings. Several factors may have contributed to differences in statistical significance, including recording environment, sample size, and the number of nights available per participant. The variation in our findings highlight the importance of examining intra-individual age effects in large samples and ensuring that participants contribute at least three nights of high-quality data when using at-home sleep EEG wearables. Furthermore, intra-individuality results from the present study must be considered preliminary, as analyses drew on data pooled from three other studies with heterogeneous recording contexts and varied sample composition. While our original findings may suggest that some sleep architecture and EEG characteristics may vary with development, future studies are necessary to validate and build upon these findings.

We note some limitations. First, the study is cross-sectional; thus, we cannot definitively state the observed age effects are due to developmental changes. Although we used linear mixed models based on exploratory visualization showing linear relationships in our sample, some developmental changes in sleep are nonlinear^26,41^, and could be better captured with flexible longitudinal analyses^65^ and larger sample sizes. Future longitudinal studies using at-home wearable sleep EEG devices will be critical for characterizing within-person developmental trajectories and addressing this limitation. Further, the current study used an automated staging algorithm validated against PSG in adults^32^, but not adolescents; our ongoing work is testing how auto-staging methods that have been trained and validated in adults perform in young people (e.g., ^66^). Although a subsample of participants completed one night of concurrent Dreem-PSG recordings, direct device validation against PSG was beyond the scope of this study, given the technical complexities involved in temporally aligning wearable device recordings with PSG sleep data. Our current work is prioritizing this comparison to further establish the validity of sleep EEG wearables in young people. Interestingly, we did not observe replicated sex differences in macro-architecture variables such as TST, SE, and WASO, though these effects have largely been derived from actigraphy and self-report^67–69^ and these measures may be more likely to reveal these effects.

This study also has several strengths. We replicated significant associations between age and sleep macro- and micro-architectural features, suggesting these effects are robust to naturalistic variation in sleep timing and duration. Our study also has strong ecological validity: participants slept in their own homes most nights, which increases generalizability to naturalistic sleep and captures habitual sleep patterns. Considering that childhood through young adulthood is a time of dramatic sleep and circadian changes, wearable sleep EEG offers the ability to capture these nuanced changes while reducing participant burden. Wearables offer feasibility and scalability, being more cost effective and accessible than lab-based PSG and enabling recruitment of larger, more diverse populations. Wearable devices also allow for longitudinal monitoring as they can be used over prolonged periods, allowing for long-term trend assessment. Taken together, these results illustrate that at-home wearable sleep devices can be used to capture age effects in sleep physiology across adolescent development.

In sum, the study leveraged sleep EEG-based wearable technology to examine age-related associations in adolescent sleep outside of the lab environment. Wearable technology offers an ecologically valid, accessible, and scalable approach to assessing developmental trajectories in sleep. Delineating normative models of sleep macro- and micro-architecture over adolescence with scalable sleep EEG devices can provide an essential template for developmentally informed sleep risk assessment and early interventions. To establish normative sleep physiology growth charts, future work would benefit from larger and more diverse samples. Our results highlight the promise of wearable sleep EEG wearable devices as a scalable tool for developmental sleep research and potential clinical assessment.

## METHODS

### Participants

All participants were recruited as part of a pilot study to demonstrate the feasibility of wearable sleep EEG data collection in young people. The pilot study was appended to three existing parent protocols across the University of Pittsburgh and Boston Children’s Hospital.

#### University of Pittsburgh (PITT)-Study 1

Twenty-three healthy adolescents (13-16 years old) were enrolled in a Dreem pilot study at the University of Pittsburgh. Participants were recruited from a larger parent protocol (P50DA046346) focused on understanding relationships between adolescent sleep-circadian rhythms and neurobehavioral mechanisms of substance use risk. Participants were eligible if they were aged 13-15 years at time of recruitment and currently enrolled in in-person schooling. Participants were excluded if they had a history of regular alcohol, cannabis, or illicit drug use in the past month; serious medical, psychiatric or neurological disorder; medication use impacting sleep/wake function (SSRIs/SSNIs excepted); MRI contraindications; sleep disorder (insomnia and delayed sleep-wake phase disorder excepted); weight <80LBS or BMI>35; or had traveled across three time zones in the past month. The University of Pittsburgh Human Rights Protection Office (HRPO) approved the study. Researchers recruited participants through a local participant registry, community advertisements, newsletters, and social media posts.

#### University of Pittsburgh (PITT)-Study 2

Forty-five adolescents and young adults (16-24 years old) were enrolled in a second Dreem pilot study at the University of Pittsburgh. Participants were recruited from a larger parent protocol focused on understanding relationships between sleep-circadian rhythms and neurobehavioral mechanisms of mood disorder risk (R01MH124828). Participants were included if they were aged 16-24 years; right handed; fluent in English; IQ>70; had no lifetime bipolar spectrum or psychosis spectrum disorder; had no mood stabilizer or antipsychotic medication use and/or changes in the previous two months; had no substance or alcohol use disorder or illicit substance use in the previous three months; no history of head trauma, neurological disorder, sleep disorder, or systemic medical illness; no MRI contraindications; not pregnant or breastfeeding; and did not engage in shift work or had extreme sleep schedules. The University of Pittsburgh HRPO approved this study, which used recruitment methods that were similar to those in PITT-Study 1.

#### Boston Children’s Hospital (BCH)

Forty individuals (9-26 years old) were recruited for a Dreem pilot study at Boston Children’s Hospital. Participants were recruited via flyers, previous study participation, the Precision Link Biobank^70^ for Healthy Discovery registries, and online posts. Inclusion criteria were age 9-26 years, fluent in English, and able to engage in research procedures. Furthermore, there could be no history of any major mental disorder (except attention deficit-hyperactivity disorder or past episode of depression), no severe neurodevelopmental disorder, brain infection, neurodegenerative disorder, or sleep disorder, no lifetime history of psychotropic medication use, no history of serious head injury, or any other serious medical condition that would make study procedures unfeasible, as assessed via self-report. The Boston Children’s Hospital Institutional Review Board approved all study procedures.

#### Analytic Sample

The combined sample comprised 108 participants enrolled across the three studies. Five participants at the PITT site were excluded due to non-viable data, and an additional three participants at the PITT site were excluded due to having less than three hours of total sleep time. All three studies followed similar data collection protocols and recruited participants from similar populations; thus, we combined data across sites to maximize our ability to detect age effects in the sample. We included a total of 100 participants for the final analysis.

### Protocol

After consent (age ≥18 years) or caregiver consent and youth assent (age <18 years) was obtained, participants completed study-specific eligibility evaluation procedures and a survey battery. We then instructed participants how to use the Dreem3 headband and complete daily electronic sleep diaries. In PITT-Study 1 and BCH, participants completed at least 3-4 consecutive nights of overnight Dreem sleep monitoring at self-selected habitual sleep times at home. In PITT-Study 2, participants completed at least 3-4 consecutive nights of Dreem monitoring: one in lab, two at-home, and then one final night of concurrent Dreem-PSG recording. We used all available nights in the present analysis. Data were collected across weekdays and weekends, spanning both the summer and academic year. Participants completed additional nights at home if needed (total nights range 1-9). Participants in PITT-1 and PITT-2 wore actigraphs. Pulse oximeters were not used in any of the three studies.

### Sleep Measures

#### Sleep EEG (Dreem3 Headband)

The Dreem3 device is a wireless headband worn during sleep that records, stores, and analyzes physiological data without requiring an active wireless connection. After data collection, the device connects via Bluetooth to a smartphone or tablet to transfer summarized metrics to a companion app, and uses Wi-Fi to transfer raw data to the sponsor’s servers. The device contains five EEG sensors (all sampled at 250 Hz, 0.4-35 Hz bandpass filter): two frontal sensors (F7, F8), two occipital sensors (O1, O2), and one ground sensor on the frontal band (FpZ location). Additional details can be found elsewhere^32^.

### Data Processing

#### Sleep Staging

The Dreem-automated algorithm performed American Academy of Sleep Medicine (AASM) sleep stage classification, which researchers have validated against PSG and manual sleep staging by expert technicians in healthy adult participants^32^. In brief, the Dreem sleep staging algorithm uses a combination of features derived from EEG, pulse oximeter, and accelerometer signals to determine the probability that each epoch belongs to each sleep stage. To predict the sleep stage for a given epoch, the algorithm incorporates features extracted from the current and past 30 epochs. In a validation study, the Dreem sleep staging algorithm correctly classified sleep stages at a level equivalent to manual sleep scoring by experts (83.8%)^32^. We examined total sleep time (TST), wake after sleep onset (WASO), and percent time spent in N1 (N1%), N2 (N2%), N3 (N3%), and REM (REM%), REM latency (time from sleep onset to the first REM epoch), time in bed (TIB), sleep efficiency (SE; TST/TIB), stage transition index from NREM to REM (TI-S), total sleep cycles, total NREM-REM cycle duration, total NREM cycle duration, and total REM cycle duration.

#### EEG preprocessing

We processed sleep EEG data using the open-source package *Luna* (https://zzz.nyspi.org/luna/). Sleep EEG was sampled at 250 Hz and band-pass filtered (0.3Hz, 35Hz) in the original records. Primary analysis focused on F7-O2 and F8-O1 channels. We first used *Luna* to convert all data to μV. Within N2, N3, and REM epochs (based on Dreem headband automated staging), we identified and removed epochs with artifacts based on the following criteria: a) delta power greater than 2.5 times the local average or beta power more than 2.0 times the local average, identified using Welch’s method, b) maximum amplitudes over 200 uV for more than 5% of the epoch, c) flat or clipped signals for more than 5% of the epoch, and d) signals 3.5 standard deviations from the mean for any of the three Hjorth parameters (i.e., activity, mobility, and complexity)^71^. We performed outlier removal based on Hjorth parameters twice for each record, following prior work^18^. We excluded channels if >50% of the epochs contained identified artifacts. In addition, we excluded records if there was <180 min of total sleep time (TST) because such records are likely to reflect unique circumstances and not typical sleep. After artifact rejection, we also used *Luna* to perform power spectral analysis, spindle detection, and slow oscillation.

#### Spectral power estimation

We used Welch’s method to estimate spectral power for NREM (N2, N3) and REM separately. For each 30-second epoch, we applied a Fast Fourier Transform using 4-second segments (0.25 spectral resolution) windowing with a Tukey taper (50%); consecutive segments overlapped by 50% (2 seconds). We summarized spectral power in the traditional frequency bands: delta (1-4Hz), theta (4-8 Hz), alpha (8-12 Hz), sigma (12-15 Hz), and beta (15-30 Hz). For each channel (F8-O1, F7-O2), we computed average power in each band for each epoch by taking 30-seconds of signal, and using Welch’s method of overlapping windows (i.e., 4-second windows within 30-second epochs), then across NREM (N2, N3) epochs separately and REM epochs separately. Relative power for each band was computed with respect to the total absolute power.

#### Automated spindle detection

We used *Luna* for automated sleep spindle detection, which relies on a validated wavelet transformation^72,73^. We detected spindles in the “slow” (11 Hz) and “fast” (15 Hz) frequencies (±2 Hz), and computed spindle density (number of spindles per minute), amplitude (based on maximum peak-to-peak amplitude), duration (seconds), mean integrated spindle activity per spindle (average amplitude and duration of an individual’s spindles), spindle chirp (within-spindle change in oscillatory frequency), and mean spindle frequency in Hz.

#### Slow oscillation detection

Following previous work^18,74^, we identified slow oscillations based on the EEG signals band-pass filtered between 0.5 and 4 Hz, focusing only on slow oscillations occurring in NREM sleep. We used the following temporal criteria to define slow oscillations: 1) a consecutive zero-crossing leading to negative peak between 0.5 and 1.5 seconds; and 2) a zero-crossing leading to positive peak not longer than 1 second. We used two approaches to measure slow oscillation amplitude: 1) an adaptive/relative threshold required that negative peak and peak-to-peak amplitudes be greater than twice the mean (for that individual/channel) and 2) an absolute threshold required a negative peak amplitude larger than -40 μV, and peak-to-peak amplitude larger than 75 μV. We estimated slow oscillation density (count per minute), mean amplitude of the negative peak, peak-to-peak amplitude, duration, and the upward slope of negative peak for each channel.

#### Slow oscillation-spindle coupling

We analyzed slow oscillation-spindle overlap for each channel and characterized coupling in *Luna*. First, we determined the proportion of spindles that coincide with a slow oscillation. Next, using the filter-Hilbert method, we calculated the slow oscillation phase at the peak of each spindle and averaged these values across slow oscillations to obtain the coupling angle for each channel. To evaluate the consistency of phase coupling between slow oscillations and spindles, we used inter-trial phase clustering to measure coupling magnitude. We z-transformed the overlap and magnitude metrics by comparing them against a null distribution generated from 10,000 random permutations, where the time series indices were shuffled^18^.

### Statistical analysis

Exploratory Locally Estimated Scatterplot Smoothing (LOESS) plots suggested most variables displayed linear associations with age in our sample. Thus, to estimate age-related changes in sleep macro- and micro-architecture, we used robust linear mixed models using the *robustlmm* package (v3.4.2)^75^ in R (v4.4.1). Because prior research indicates that sleep physiology may differ between males and females^13,76^, we also examined sex effects. We included age and sex as fixed effects, and specified random intercepts for participants nested within site to account for clustering by site and repeated measures across nights. For microarchitecture variables, we also specified channel (F7 or F8) as a fixed effect. We explored including an autoregressive correlation structure across nights which yielded very similar estimates; therefore, all reported results are from the simpler model with random intercepts for subjects nested within site. We used eight macro- and fifteen micro-architecture measures as our primary outcome variables. We used False Discovery Rate (FDR) to correct for multiple comparisons (N=23), and corrected p-values (*p*_FDR_) < 0.05 to define statistical significance^77^. Sleep characteristics can vary between weeknights and weekends and across the summer and school year due to social schedules, circadian misalignment, and recovery from prior sleep restriction^78–80^. Therefore, to further explore potential confounds, we also ran post-hoc models that included weekday/weekend night status and summer vs. school year as fixed effects in the primary model.

We first examined sleep macro-architecture (N=8 variables) and micro-architecture measures (N=15 variables) with known age effects from traditional PSG (i.e., at least two studies with consistent age effects; Table S1 & S2). We next explored macro-architecture variables with lesser-known age effects (i.e., unexplored, less than two studies with consistent age-effects; N=5 variables): stage transition index from NREM to REM (TI-S), total number of sleep cycles, total cycle duration, total NREM cycle duration, and total REM cycle duration. We further explored micro-architecture with lesser-known age effects (N=19 variables): slow oscillation duration, slow oscillation peak-to-peak amplitude, relative NREM delta, sigma, theta, and beta power, relative REM delta, sigma, theta, beta, alpha power, and absolute REM alpha and beta power; integrated slow spindle activity, integrated fast spindle activity, slow and fast spindle chirp, and slow and fast spindle frequency.

#### Intra-individual variability

For participants with at least three nights of data (N=80, 80% of the sample), we calculated intra-individual variability for all macro- and micro-architecture variables. To enable comparison with previous studies^56,58,59,81,82^, we used a commonly used measure of intra-individual variability, the individual standard deviation^83^. We ran robust linear models using the *lmrob* function from the *robust* R package^84^, covarying for age, sex and site. Our outcome variable was individual standard deviation across nights for all variables. We corrected for multiple comparisons across measures (N=47) using the False Discovery Rate^77^.

## DATA AVAILABILITY

The datasets analyzed in the current study are not publicly available due to privacy and confidentiality laws but summarized data are available from the corresponding author on reasonable request.

## CODE AVAILABILITY

The underlying code for this study will be made available upon publication.

## ACKNOWLEDGMENTS

The authors thank all participants and families in this study. They also thank all research assistants who contributed to data collection.

This work was supported by the National Institute of Mental Health grants R01MH129636 (Jalbrzikowski), R01MH124828 (Soehner), the National Heart Lung and Blood Institute grant R01HL169318 (Soehner, Jalbrzikowski) and the National Institute on Drug Abuse grant P50DA046346 (Franzen). MJ was also supported by the Tommy Fuss Center for Neuropsychiatric Research Next Generation Award. SL was supported by the National Research Service Award (T32MH112510). The funders played no role in study design, data collection, analysis and interpretation of data, or the writing of this manuscript.

This work was previously published on *bioRxiv*. https://doi.org/10.1101/2025.10.10.681690

## AUTHOR CONTRIBUTIONS

**SL:** Methodology, Formal analysis, Investigation, Writing – Original Draft, Writing – Review & Editing, Visualization. **RC:** Formal analysis, Investigation, Writing – Original Draft, Writing – Review & Editing, Visualization. **RH:** Data curation. **SS:** Investigation, Writing – Review & Editing. **IE:** Investigation, Writing – Review & Editing**. MC:** Investigation, Writing – Review & Editing**. BH:** Investigation, Writing – Review & Editing**. MF-W:** Investigation, Writing – Review & Editing. **LK:** Investigation, Writing – Review & Editing. **SC:** Data curation. **PF:** Writing – Review & Editing, Funding acquisition. **DB:** Writing – Review & Editing. **BPH:** Writing – Review & Editing. **JL:** Writing – Review & Editing. **MLW:** Writing – Review & Editing. **DBC:** Writing – Review & Editing. **RGB:** Writing – Review & Editing. **AS:** Conceptualization, Methodology, Writing – Review & Editing, Supervision, Project administration, Funding acquisition. **MJ:** Conceptualization, Methodology, Writing – Review & Editing, Supervision, Project administration, Funding acquisition.

