## Supplemental for "Using wearable EEG to examine age trends in sleep macro- and micro-architecture across adolescence"

**Supplement.**

| Table S1. Studies assessing the age effects of sleep macro-architecture | 2 |
| --- | --- |
| Table S2. Studies assessing the age-effects of sleep micro-architecture | 3 |
| Table S3. Mean, SD, and range for established sleep macro- and micro-architecture variables | 4 |
| Table S4. Effect of age on sleep macro-architecture across adolescence for at-home recordings only | 5 |
| Table S5. Effect of age on sleep micro-architecture across adolescence for at-home recordings only | 6 |
| Table S6. Effect of age on sleep macro-architecture across adolescence when accounting for  weekday/weeknight night of recording | 7 |
| Table S7. Effect of age on sleep macro-architecture across adolescence when accounting  for summer/school year night of recording | 8 |
| Table S8. Effect of age on sleep micro-architecture across adolescence when accounting for  weekday/weekend night of recording | 9 |
| Table S9. Effect of age on sleep micro-architecture across adolescence when accounting for summer/school year night of recording | 10 |
| Table S10. Effect of sex on sleep macro- and microarchitecture variables | 11 |
| Table S11. Effect of age on intra-individual variability of macro- and microarchitecture variables | 13 |
| Table S12. Effect of age on intra-individual variability of macro- and microarchitecture variables for at-home recordings only  Table S13. Within-subjects comparisons of macro- and microarchitecture variables between at-home and in-lab recordings. | 15  17 |
| Figure S1. Significant associations between sleep macro-architecture and age, separated by sex. | 19 |
| Figure S2. Associations between sleep micro-architecture and age | 20 |
| Figure S3. Macro- and micro-architecture variables showing significant sex differences | 21 |
| Figure S4. Age-related associations in micro-architecture variables not replicated in the present study | 22 |
| Figure S5. Significant associations between age and intra-individual variability of macro- and micro-architecture variables | 23 |
| Supplemental References | 24 |

Table S1. Electroencephalography studies assessing the effect of age on sleep macro-architecture.

| **Variable** | **Reference #1** | **Reference #1 Age(s) Tested** | **Reference #2** | **Reference #2 Age(s) Tested** | **Age effect** |
| --- | --- | --- | --- | --- | --- |
| Total Sleep Time (TST) | Kozhemiako et al., 2024^1^ | 2.5-17.5 years | Scholle et al., 2011^2^ | 1-18 years | TST decreases with age |
| Wake After Sleep Onset (WASO) | Kozhemiako et al., 2024^1^ | 2.5-17.5 years | Baker et al., 2016^3^ | 12-21 years | WASO increases with age |
| N2% | Kozhemiako et al., 2024^1^ | 2.5-17.5 years | Scholle et al., 2011^2^ | 1-18 years | N2% increases with age |
| N3% | Tarokh & Carskadon, 2010^4^ | 9-13 years | Baker et al., 2016^3^ | 12-21 years | N3% decreases with age |
| Rapid Eye-Movement (REM) sleep (%) | Jenni & Carskadon, 2004^5^ | 9-16 years | Kozhemiako et al., 2024^1^ | 2.5-17.5 years | REM% decreases with age |
| REM Latency | Coble et al., 1984^6^ | 6-16 years | Gillin et al., 1981^7^ | 15-64 years | REM latency decreases with age |
| Sleep Efficiency | Kozhemiako et al., 2024^1^ | 2.5-17.5 years | Baker et al., 2016^3^ | 12-21 years | Sleep efficiency decreases with age |
| Time In Bed (TIB) | Feinberg et al., 2012^8^ | 9-18 years | Coble et al., 1984^6^ | 6-16 years | TIB decreases with age |

Table S2. Electroencephalography studies assessing the effect of age on sleep micro-architecture.

| **Variable** | **Reference #1** | **Reference #1 Age(s) Tested** | **Reference #2** | **Reference #2 Age(s) Tested** | **Age effect** |
| --- | --- | --- | --- | --- | --- |
| Absolute NREM delta power | Baker et al., 2016^3^ | 12-21 years | Tarokh & Carskadon, 2010^4^ | 9-13 years | NREM delta power decreases with age |
| Absolute NREM sigma power | Goldstone et al., 2019^10^ | 12-21 years | Kurth et al., 2010^11^ | 2.4-19.4 years | NREM sigma power increases until age 11 then decreases with age |
| Absolute NREM theta power | Campbell & Feinberg, 2009^12^ | 6-17 years | Campbell et al., 2012^13^ | 9-18 years | NREM theta decreases with age |
| Absolute NREM beta power | Tarokh & Carskadon, 2010^4^ | 9-13 years | Gaudreau, Carrier & Montplaisir, 2001^14^ | 6-60 years | NREM beta decreases with age |
| Absolute REM delta power | Baker et al., 2012^15^ | 11-14 years | Feinberg & Campbell, 2013^16^ | 6-18 years | REM delta power decreases with age |
| Absolute REM sigma power | Jenni & Carskadon, 2004^5^ | 9-16 years | Baker et al., 2012^15^ | 11-14 years | REM sigma power decreases with age |
| Absolute REM theta power | Tarokh & Carskadon, 2010^4^ | 9-13 years | Baker et al., 2012^15^ | 11-14 years | REM theta power decreases with age |
| Slow oscillation (SO)-spindle coupling | Joechner et al., 2023^17^ | 526 years | Hahn et al., 2020^18^ | 9.5-16 years (mean ages) | SO-spindle coupling is temporally more precise with age |
| Slow spindle amplitude | Goldstone et al., 2019^10^ | 12-21 years | Nader & Smith, 2015^19^ | 12-19 years | Slow spindle amplitude decreases with age |
| Slow spindle density | Gombos et al., 2022^20^ | 12-20 years | Nader & Smith, 2015^19^ | 12-19 years | Slow spindle density decreases with age |
| Slow spindle duration | Gombos et al., 2022^20^ | 12.5-21.3 years (mean ages) | Goldstone et al., 2019^10^ | 12-21 years | Slow spindle duration decreases with age |
| Fast spindle amplitude | Goldstone et al., 2019^10^ | 12-21 years | Nader & Smith, 2015^19^ | 12-19 years | Fast spindle amplitude decreases with age |
| Fast spindle density | Goldstone et al., 2019^10^ | 12-21 years | Nader & Smith, 2015^19^ | 12-19 years | Fast spindle density increases with age |
| Fast spindle duration | Gombos et al., 2022^20^ | 12.5-21.3 years (mean ages) | Goldstone et al., 2019^10^ | 12-21 years | Fast spindle duration decreases with age |

Abbreviations: NREM, non-rapid eye movement sleep; REM, rapid eye movement sleep.

Table S3. Mean, SD, and range for established sleep macro- and micro-architecture variables.

|  | **Total (N = 100)** | | | **BCH (N = 40)** | | | **PITT (N = 60)** | | |
| --- | --- | --- | --- | --- | --- | --- | --- | --- | --- |
| **Sleep Variables** | **Mean** | **SD** | **Range** | **Mean** | **SD** | **Range** | **Mean** | **SD** | **Range** |
| Total Sleep Time (min) | 408.59 | 87.74 | 181-622.5 | 426.51 | 89.48 | 203-622.5 | 397.58 | 85.02 | 181-619 |
| Wake After Sleep Onset | 31.25 | 38.18 | 2-249.5 | 26.50 | 27.69 | 3.5-133.5 | 34.17 | 43.20 | 2-249.5 |
| N2% | 41.48 | 8.86 | 14.5-65.97 | 39.31 | 8.42 | 18.73-59.78 | 42.81 | 8.88 | 14.5-65.97 |
| N3% | 28.76 | 9.51 | 5.69-61.36 | 30.77 | 9.39 | 11.26-50.88 | 27.53 | 9.40 | 5.69-61.36 |
| REM% | 24.44 | 8.31 | 1.37-74.93 | 24.84 | 7.21 | 5.91-53.16 | 24.19 | 8.93 | 1.37-74.93 |
| REM Latency | 97.54 | 49.48 | 0-296 | 98.72 | 44.54 | 0-218.5 | 96.81 | 52.38 | 2-296 |
| Sleep Efficiency | 85.72 | 13.68 | 12.67-98.66 | 87.60 | 11.51 | 36.38-97.87 | 84.56 | 14.77 | 12.67-98.66 |
| Time in Bed | 474.69 | 92.72 | 209.5-880.5 | 488.83 | 91.91 | 271-880.5 | 465.99 | 92.39 | 209.5-862 |
| Absolute NREM Delta | 905.34 | 440.58 | 181.66-2375.54 | 940.60 | 461.07 | 181.66-2189.05 | 883.67 | 427.28 | 242.86-2375.54 |
| Absolute NREM Sigma | 11.99 | 8.17 | 2.88-83.81 | 10.50 | 9.09 | 2.88-83.81 | 12.91 | 7.43 | 3.28-48.59 |
| Absolute NREM Theta | 111.14 | 83.99 | 21.97-784.65 | 101.61 | 73.74 | 21.97-574.4 | 117.01 | 89.40 | 25.45-784.65 |
| Absolute NREM Beta | 12.83 | 16.20 | 3.41-195.12 | 12.87 | 22.77 | 3.41-195.12 | 12.80 | 10.35 | 4.5-92.27 |
| Absolute REM Delta | 404.05 | 416.63 | 30.69-3380.83 | 328.31 | 319.93 | 56.75-1581.22 | 450.60 | 460.86 | 30.69-3380.83 |
| Absolute REM Sigma | 9.77 | 12.85 | 1.81-125.4 | 6.65 | 5.68 | 1.81-34.2 | 11.68 | 15.41 | 2.14-125.4 |
| Absolute REM Theta | 102.85 | 117.99 | 12.35-779.64 | 78.40 | 99.96 | 12.78-627.38 | 117.88 | 125.71 | 12.35-779.64 |
| Slow spindle duration | 0.87 | 0.11 | 0.67-1.33 | 0.91 | 0.13 | 0.72-1.33 | 0.84 | 0.09 | 0.67-1.29 |
| Slow spindle amplitude | 45.42 | 14.98 | 21.28-102.79 | 43.39 | 14.46 | 21.28-102.49 | 46.66 | 15.20 | 21.62-102.79 |
| Slow spindle density | 1.29 | 0.63 | 0.01-3.56 | 1.52 | 0.73 | 0.52-3.56 | 1.15 | 0.50 | 0.01-2.65 |
| Fast spindle duration | 0.78 | 0.08 | 0.59-1.17 | 0.80 | 0.08 | 0.68-1.11 | 0.76 | 0.08 | 0.59-1.17 |
| Fast spindle amplitude | 28.84 | 11.69 | 13.02-88.3 | 28.74 | 13.48 | 13.5-88.3 | 28.90 | 10.48 | 13.02-74.97 |
| Fast spindle density | 0.48 | 0.33 | 0.02-1.89 | 0.50 | 0.33 | 0.1-1.72 | 0.47 | 0.32 | 0.02-1.89 |
| Slow oscillation-slow spindle coupling magnitude | 0.21 | 0.11 | 0.02-1 | 0.17 | 0.07 | 0.04-0.36 | 0.23 | 0.12 | 0.02-1 |
| Slow oscillation-fast spindle coupling magnitude | 0.18 | 0.12 | 0.03-0.74 | 0.15 | 0.09 | 0.03-0.52 | 0.20 | 0.13 | 0.04-0.74 |

Abbreviations: NREM, non-rapid eye movement sleep; REM, rapid eye movement slee

**Table S4. Effect of age on sleep macro-architecture across adolescence using at-home**

**recordings only.**

| Sleep Macro-architecture | | | | | |
| --- | --- | --- | --- | --- | --- |
| Sleep Variables | β | SE | t | *p* | *p*_FDR_ |
| Total Sleep Time | -0.13 | 0.08 | -1.65 | 9.97e-02 | 1.14e-01 |
| Wake After Sleep Onset | -0.06 | 0.04 | -1.45 | 1.47e-01 | 1.47e-01 |
| N2% | **0.23** | **0.09** | **2.71** | **6.95e-03** | **1.39e-02** |
| N3% | **-0.43** | **0.08** | **-5.21** | **2.84e-07** | **2.27e-06** |
| REM% | **0.16** | **0.06** | **2.48** | **1.34e-02** | **2.14e-02** |
| REM latency | **-0.25** | **0.07** | **-3.83** | **1.46e-04** | **3.89e-04** |
| Sleep Efficiency | 0.09 | 0.06 | 1.72 | 8.62e-02 | 1.14e-01 |
| Time In Bed | **-0.27** | **0.07** | **-3.90** | **1.09e-04** | **3.89e-04** |

Abbreviations: N, non-rapid eye movement sleep; REM, rapid eye movement sleep. Models

account for age, sex, and weekday/weekend night of recording as fixed effects, and participant

nested within site as a random effect. Values in bold survive correction for multiple comparisons

(false discovery rate, *p*_FDR_<0.05).

**Table S5. Effect of age on sleep micro-architecture across adolescence using at-home recordings only.**

|  | **Sleep Micro-architecture** | | | | | |
| --- | --- | --- | --- | --- | --- | --- |
| Absolute power during NREM and REM | Sleep Variables | β | SE | t | *p* | *p*_FDR_ |
|  | NREM Delta | **-0.50** | **0.07** | **-7.33** | **1.05e-12** | **7.84e-12** |
|  | NREM Sigma | -0.02 | 0.05 | -0.47 | 6.38e-01 | 6.83e-01 |
|  | NREM Theta | **-0.12** | **0.04** | **-2.68** | **7.54e-03** | **1.61e-02** |
|  | NREM Beta | -0.01 | 0.02 | -0.67 | 5.05e-01 | 6.49e-01 |
|  | REM Delta | -0.03 | 0.04 | -0.65 | 5.19e-01 | 6.49e-01 |
|  | REM Sigma | 0.02 | 0.02 | 0.79 | 4.32e-01 | 6.48e-01 |
|  | REM Theta | 0.00 | 0.03 | 0.02 | 9.85e-01 | 9.85e-01 |
| Sleep spindle and coupling features | Slow spindle duration | **-0.43** | **0.06** | **-7.50** | **3.33e-13** | **5.00e-12** |
|  | Slow spindle amplitude | -0.12 | 0.06 | -2.02 | 4.44e-02 | 8.32e-02 |
|  | Slow spindle density | **-0.43** | **0.08** | **-4.95** | **1.07e-06** | **5.36e-06** |
|  | Fast spindle duration | **-0.18** | **0.04** | **-4.21** | **3.09e-05** | **1.16e-04** |
|  | Fast spindle amplitude | -0.03 | 0.05 | -0.57 | 5.68e-01 | 6.56e-01 |
|  | Fast spindle density | -0.07 | 0.06 | -1.20 | 2.31e-01 | 3.85e-01 |
|  | Slow oscillation-slow spindle coupling magnitude | **0.14** | **0.04** | **3.25** | **1.22e-03** | **3.66e-03** |
|  | Slow oscillation-fast spindle coupling magnitude | **0.11** | **0.04** | **3.06** | **2.31e-03** | **5.79e-03** |

Abbreviations: NREM, non-rapid eye movement sleep; REM, rapid eye movement sleep. Models account for age, sex and channel as fixed effects, and participant nested within site as a random effect. Values in bold survive correction for multiple comparisons (false discovery rate, *p*_FDR_<0.05)

Table S6. Effect of age on sleep macro-architecture across adolescence when accounting for weekday/weekend night of recording.

| Sleep Macro-architecture | | | | | |
| --- | --- | --- | --- | --- | --- |
| Sleep Variables | β | SE | t | *p* | *p*_FDR_ |
| Total Sleep Time | -0.14 | 0.07 | -1.88 | 6.04e-02 | 9.66e-02 |
| Wake After Sleep Onset | -0.06 | 0.04 | -1.33 | 1.85e-01 | 1.85e-01 |
| N2% | **0.28** | **0.08** | **3.50** | **5.04e-04** | **1.01e-03** |
| N3% | **-0.39** | **0.08** | **-4.88** | **1.38e-06** | **1.10e-05** |
| REM% | 0.11 | 0.07 | 1.62 | 1.07e-01 | 1.32e-01 |
| REM latency | **-0.24** | **0.06** | **-3.73** | **2.11e-04** | **8.42e-04** |
| Sleep Efficiency | 0.08 | 0.05 | 1.58 | 1.15e-01 | 1.32e-01 |
| Time In Bed | **-0.25** | **0.07** | **-3.57** | **3.85e-04** | **1.01e-03** |

Abbreviations: N, non-rapid eye movement sleep; REM, rapid eye movement sleep. Models

account for age, sex, and weekday/weekend night of recording as fixed effects, and participant

nested within site as a random effect. Values in bold survive correction for multiple comparisons

(false discovery rate, *p*_FDR_<0.05).

Table S7. Effect of age on sleep macro-architecture across adolescence when accounting for summer/school year night of recording.

| Sleep Macro-architecture | | | | | |
| --- | --- | --- | --- | --- | --- |
| Sleep Variables | β | SE | t | *p* | *p*_FDR_ |
| Total Sleep Time | -0.14 | 0.08 | -1.85 | 6.49e-02 | 8.65e-02 |
| Wake After Sleep Onset | -0.06 | 0.04 | -1.32 | 1.86e-01 | 1.86e-01 |
| N2% | **0.25** | **0.08** | **3.13** | **1.85e-03** | **3.69e-03** |
| N3% | **-0.39** | **0.08** | **-4.67** | **3.67e-06** | **2.93e-05** |
| REM% | 0.14 | 0.07 | 1.86 | 6.28e-02 | 8.65e-02 |
| REM latency | **-0.26** | **0.07** | **-3.92** | **9.85e-05** | **3.94e-04** |
| Sleep Efficiency | 0.09 | 0.05 | 1.75 | 8.06e-02 | 9.21e-02 |
| Time In Bed | **-0.25** | **0.07** | **-3.45** | **5.96e-04** | **1.59e-03** |

Abbreviations: NREM, non-rapid eye movement sleep; REM, rapid eye movement sleep. Models

account for age, sex and summer/school year night of recording as fixed effects, and participant

nested within site as a random effect. Values in bold survive correction for multiple comparisons

(false discovery rate, *p*_FDR_<0.05).

Table S8. Effect of age on sleep micro-architecture across adolescence when accounting for weekday/weekend night of recording.

|  | Sleep Micro-architecture | | | | | |
| --- | --- | --- | --- | --- | --- | --- |
| Absolute power during NREM and REM | Sleep Variables | β | SE | t | *p* | *p*_FDR_ |
|  | NREM Delta | **-0.50** | **0.07** | **-7.33** | **8.16e-13** | **6.12e-12** |
|  | NREM Sigma | -0.04 | 0.04 | -0.85 | 3.97e-01 | 5.50e-01 |
|  | NREM Theta | **-0.11** | **0.04** | **-2.96** | **3.22e-03** | **8.05e-03** |
|  | NREM Beta | -0.01 | 0.02 | -0.71 | 4.77e-01 | 5.50e-01 |
|  | REM Delta | -0.02 | 0.03 | -0.75 | 4.54e-01 | 5.50e-01 |
|  | REM Sigma | 0.01 | 0.02 | 0.65 | 5.14e-01 | 5.50e-01 |
|  | REM Theta | 0.01 | 0.03 | 0.21 | 8.30e-01 | 8.30e-01 |
| Sleep spindle and coupling features | Slow spindle duration | **-0.41** | **0.06** | **-7.55** | **1.76e-13** | **2.63e-12** |
|  | Slow spindle amplitude | -0.11 | 0.06 | -2.00 | 4.57e-02 | 8.56e-02 |
|  | Slow spindle density | **-0.43** | **0.08** | **-5.41** | **9.23e-08** | **4.61e-07** |
|  | Fast spindle duration | **-0.17** | **0.04** | **-4.23** | **2.76e-05** | **1.03e-04** |
|  | Fast spindle amplitude | -0.04 | 0.05 | -0.78 | 4.35e-01 | 5.50e-01 |
|  | Fast spindle density | -0.06 | 0.06 | -1.10 | 2.71e-01 | 4.52e-01 |
|  | Slow oscillation-slow spindle coupling magnitude | **0.15** | **0.04** | **3.42** | **6.73e-04** | **2.02e-03** |
|  | Slow oscillation-fast spindle coupling magnitude | **0.10** | **0.04** | **2.78** | **5.67e-03** | **1.22e-02** |

Abbreviations: NREM, non-rapid eye movement sleep; REM, rapid eye movement sleep; SO, slow oscillation. Models account for age, sex, channel, and weekday/weekend night of recording as fixed effects, and participant nested within site as a random effect. Values in bold survive correction for multiple comparisons (false discovery rate, *p*_FDR_<0.05).

Table S9. Effect of age on sleep micro-architecture across adolescence when accounting for summer/school year night of recording.

| Outcome | | β | SE | t | *p* | *p*_FDR_ |
| --- | --- | --- | --- | --- | --- | --- |
| Absolute power during NREM and REM | NREM Delta | **-0.53** | **0.07** | **-7.45** | **3.42e-13** | **2.57e-12** |
|  | NREM Sigma | -0.04 | 0.04 | -0.85 | 3.97e-01 | 5.45e-01 |
|  | NREM Theta | **-0.11** | **0.04** | **-2.96** | **3.22e-03** | **8.05e-03** |
|  | NREM Beta | -0.01 | 0.02 | -0.81 | 4.18e-01 | 5.45e-01 |
|  | REM Delta | -0.02 | 0.03 | -0.75 | 4.54e-01 | 5.45e-01 |
|  | REM Sigma | 0.01 | 0.02 | 0.65 | 5.14e-01 | 5.50e-01 |
|  | REM Theta | 0.01 | 0.03 | 0.21 | 8.30e-01 | 8.30e-01 |
| Sleep spindle and coupling features | Slow spindle duration | **-0.41** | **0.06** | **-7.55** | **1.76e-13** | **2.57e-12** |
|  | Slow spindle amplitude | -0.11 | 0.06 | -2.00 | 4.57e-02 | 8.56e-02 |
|  | Slow spindle density | **-0.43** | **0.08** | **-5.41** | **9.23e-08** | **4.61e-07** |
|  | Fast spindle duration | **-0.17** | **0.04** | **-4.23** | **2.76e-05** | **1.03e-04** |
|  | Fast spindle amplitude | -0.03 | 0.05 | -0.72 | 4.73e-01 | 5.45e-01 |
|  | Fast spindle density | -0.06 | 0.06 | -0.95 | 3.42e-01 | 5.45e-01 |
|  | Slow oscillation-slow spindle coupling magnitude | **0.15** | **0.04** | **3.42** | **6.73e-04** | **2.02e-03** |
|  | Slow oscillation-fast spindle coupling magnitude | **0.10** | **0.04** | **2.78** | **5.67e-03** | **1.22e-02** |

Abbreviations: NREM, non-rapid eye movement sleep; REM, rapid eye movement sleep. Models account for age, sex, channel, and summer/school year night of recording as fixed effects, and participant nested within site as a random effect. Values in bold survive correction for multiple comparisons (false discovery rate, *p*_FDR_<0.05).

Table S10. Effect of sex on sleep macro- and microarchitecture variables.

| Outcome | | β | SE | t | *p* | *p*_FDR_ |
| --- | --- | --- | --- | --- | --- | --- |
| Sleep macro-architecture features | Total Sleep Time | -0.29 | 0.15 | -1.95 | 5.20E-02 | 2.08E-01 |
|  | Wake After Sleep Onset | -0.05 | 0.08 | -0.63 | 5.30E-01 | 8.49E-01 |
|  | N2% | -0.19 | 0.16 | -1.23 | 2.20E-01 | 5.69E-01 |
|  | N3% | 0.17 | 0.16 | 1.07 | 2.85E-01 | 5.69E-01 |
|  | REM% | 0.05 | 0.14 | 0.38 | 7.01E-01 | 9.20E-01 |
|  | REM latency | 0.01 | 0.13 | 0.10 | 9.20E-01 | 9.20E-01 |
|  | Sleep Efficiency | -0.02 | 0.10 | -0.16 | 8.76E-01 | 9.20E-01 |
|  | Time In Bed | -0.33 | 0.14 | -2.46 | 1.42E-02 | 1.14E-01 |
|  | Transition index from NREM to REM | -0.03 | 0.15 | -0.17 | 8.67E-01 | 8.67E-01 |
|  | Total sleep cycles | **-0.37** | **0.12** | **-2.96** | **3.15E-03** | **1.58E-02** |
|  | Total cycle duration | -0.28 | 0.15 | -1.90 | 5.78E-02 | 1.14E-01 |
|  | Total NREM cycle duration | -0.27 | 0.15 | -1.82 | 6.85E-02 | 1.14E-01 |
|  | Total REM cycle duration | -0.09 | 0.16 | -0.56 | 5.76E-01 | 7.19E-01 |
| Sleep spindle and coupling features | Slow spindle duration | -0.02 | 0.11 | -0.17 | 8.66E-01 | 8.66E-01 |
|  | Slow spindle amplitude | 0.02 | 0.11 | 0.21 | 8.35E-01 | 8.66E-01 |
|  | Slow spindle density | 0.08 | 0.16 | 0.48 | 6.28E-01 | 8.66E-01 |
|  | Fast spindle duration | -0.05 | 0.08 | -0.69 | 4.87E-01 | 8.66E-01 |
|  | Fast spindle amplitude | 0.03 | 0.09 | 0.37 | 7.09E-01 | 8.66E-01 |
|  | Fast spindle density | -0.17 | 0.11 | -1.45 | 1.47E-01 | 8.66E-01 |
|  | Slow oscillation-slow spindle coupling magnitude | 0.03 | 0.09 | 0.29 | 7.69E-01 | 8.66E-01 |
|  | Slow oscillation-fast spindle coupling magnitude | 0.04 | 0.07 | 0.62 | 5.34E-01 | 8.66E-01 |
| Slow oscillation features | Slow oscillation duration | 0.14 | 0.15 | 0.93 | 3.53E-01 | 5.30E-01 |
|  | Median slow oscillation peak to peak amplitude | 0.00 | 0.13 | -0.03 | 9.77E-01 | 9.77E-01 |
| Relative power during NREM and REM | NREM Delta | -0.15 | 0.18 | -0.86 | 3.92E-01 | 5.30E-01 |
|  | NREM Sigma | -0.10 | 0.11 | -0.99 | 3.23E-01 | 5.30E-01 |
|  | NREM Theta | -0.06 | 0.11 | -0.56 | 5.73E-01 | 6.81E-01 |
|  | NREM Beta | -0.06 | 0.04 | -1.31 | 1.92E-01 | 4.05E-01 |
|  | REM Delta | -0.11 | 0.13 | -0.86 | 3.91E-01 | 5.30E-01 |
|  | REM Sigma | **0.36** | **0.11** | **3.17** | **1.59E-03** | **1.51E-02** |
|  | REM Theta | **0.34** | **0.10** | **3.36** | **8.40E-04** | **1.51E-02** |
|  | REM Alpha | 0.21 | 0.11 | 1.87 | 6.21E-02 | 1.69E-01 |
|  | REM Beta | **0.17** | **0.06** | **2.90** | **3.89E-03** | **2.46E-02** |
| Absolute power during NREM and REM | NREM Delta | -0.08 | 0.13 | -0.60 | 5.50E-01 | 8.66E-01 |
|  | NREM Sigma | -0.07 | 0.08 | -0.82 | 4.10E-01 | 8.66E-01 |
|  | NREM Theta | -0.03 | 0.07 | -0.44 | 6.59E-01 | 8.66E-01 |
|  | NREM Beta | -0.02 | 0.03 | -0.51 | 6.10E-01 | 8.66E-01 |
|  | REM Delta | -0.09 | 0.06 | -1.46 | 1.45E-01 | 8.66E-01 |
|  | REM Sigma | -0.01 | 0.04 | -0.27 | 7.86E-01 | 8.66E-01 |
|  | REM Theta | -0.04 | 0.05 | -0.81 | 4.19E-01 | 8.66E-01 |
|  | REM Alpha | -0.05 | 0.05 | -1.19 | 2.34E-01 | 4.46E-01 |
|  | REM Beta | 0.00 | 0.01 | -0.25 | 8.04E-01 | 8.49E-01 |
| Sleep spindle features | Integrated slow spindle activity | -0.06 | 0.12 | -0.45 | 6.52E-01 | 7.28E-01 |
|  | Integrated fast spindle activity | -0.22 | 0.10 | -2.26 | 2.43E-02 | 1.15E-01 |
|  | Slow spindle chirp | 0.28 | 0.17 | 1.71 | 8.85E-02 | 2.10E-01 |
|  | Fast spindle chirp | 0.25 | 0.12 | 2.02 | 4.37E-02 | 1.38E-01 |
|  | Slow spindle frequency | -0.37 | 0.17 | -2.11 | 3.54E-02 | 1.34E-01 |
|  | Fast spindle frequency | 0.14 | 0.17 | 0.81 | 4.19E-01 | 5.30E-01 |

Abbreviations: NREM, non-rapid eye movement sleep; REM, rapid eye movement sleep. Models account for age and sex as fixed effects, and participant nested within site as a random effect. Microarchitecture variables also included channel as fixed effect. Values in bold survive correction for multiple comparisons (false discovery rate, *p*_FDR_<0.05).

Table S11. Effect of age on intra-individual variability of macro- and microarchitecture variables.

| Outcome | | β | SE | t | *p* | *p*_FDR_ |
| --- | --- | --- | --- | --- | --- | --- |
| Sleep macro-architecture features | Total Sleep Time | 0.01 | 0.12 | 0.09 | 9.28e-01 | 9.45e-01 |
|  | Wake After Sleep Onset | -0.12 | 0.08 | -1.56 | 1.23e-01 | 2.47e-01 |
|  | N2% | -0.01 | 0.13 | -0.07 | 9.45e-01 | 9.45e-01 |
|  | N3% | 0.10 | 0.11 | 0.88 | 3.80e-01 | 5.07e-01 |
|  | REM% | **0.21** | **0.06** | **3.33** | **1.35e-03** | **1.08e-02** |
|  | REM latency | -0.24 | 0.11 | -2.07 | 4.21e-02 | 1.68e-01 |
|  | Sleep Efficiency | -0.10 | 0.09 | -1.08 | 2.82e-01 | 4.52e-01 |
|  | Time In Bed | 0.12 | 0.07 | 1.71 | 9.15e-02 | 2.44e-01 |
|  | Transition index from NREM to REM | 0.06 | 0.11 | 0.51 | 6.13e-01 | 9.23e-01 |
|  | Total sleep cycles | -0.16 | 0.10 | -1.62 | 1.09e-01 | 5.44e-01 |
|  | Total cycle duration | -0.02 | 0.13 | -0.18 | 8.57e-01 | 9.23e-01 |
|  | Total NREM cycle duration | -0.01 | 0.12 | -0.10 | 9.23e-01 | 9.23e-01 |
|  | Total REM cycle duration | 0.07 | 0.12 | 0.57 | 5.68e-01 | 9.23e-01 |
| Sleep spindle and coupling features | Slow spindle duration | -0.03 | 0.07 | -0.46 | 6.44e-01 | 8.20e-01 |
|  | Slow spindle amplitude | 0.12 | 0.11 | 1.12 | 2.66e-01 | 7.15e-01 |
|  | Slow spindle density | 0.07 | 0.09 | -0.83 | 4.07e-01 | 7.70e-01 |
|  | Fast spindle duration | -0.02 | 0.10 | -0.23 | 8.21e-01 | 9.32e-01 |
|  | Fast spindle amplitude | 0.06 | 0.07 | 0.84 | 4.06e-01 | 7.15e-01 |
|  | Fast spindle density | 0.03 | 0.06 | 0.49 | 6.22e-01 | 8.20e-01 |
|  | Slow oscillation-slow spindle coupling magnitude | **0.25** | **0.08** | **3.16** | **2.28e-03** | **3.20e-02** |
|  | Slow oscillation-fast spindle coupling magnitude | 0.18 | 0.12 | 1.49 | 1.41e-01 | 6.57e-01 |
| Slow oscillation features | Slow oscillation duration | 0.18 | 0.08 | 2.27 | 2.61e-02 | 9.90e-02 |
|  | Median slow oscillation peak to peak amplitude | -0.03 | 0.08 | -0.45 | 6.55e-01 | 7.32e-01 |
| Relative power during NREM and REM | NREM Delta | 0.16 | 0.11 | 1.48 | 1.44e-01 | 3.04e-01 |
|  | NREM Sigma | 0.17 | 0.06 | 2.61 | 1.08e-02 | 5.11e-02 |
|  | NREM Theta | **0.17** | **0.06** | **2.93** | **4.41e-03** | **3.85e-02** |
|  | NREM Beta | **0.03** | **0.01** | **3.07** | **3.00e-03** | **3.85e-02** |
|  | REM Delta | 0.19 | 0.14 | 1.34 | 1.83e-01 | 3.06e-01 |
|  | REM Sigma | 0.10 | 0.05 | 1.98 | 5.18e-02 | 1.41e-01 |
|  | REM Theta | 0.11 | 0.09 | 1.22 | 2.25e-01 | 3.06e-01 |
|  | REM Alpha | 0.11 | 0.05 | 2.15 | 3.46e-02 | 1.10e-01 |
|  | REM Beta | 0.02 | 0.01 | 1.23 | 2.23e-01 | 3.06e-01 |
| Absolute power during NREM and REM | NREM Delta | -0.22 | 0.08 | -2.70 | 8.57e-03 | 6.00e-02 |
|  | NREM Sigma | 0.03 | 0.04 | 0.87 | 3.84e-01 | 7.15e-01 |
|  | NREM Theta | 0.00 | 0.06 | 0.09 | 9.32e-01 | 9.32e-01 |
|  | NREM Beta | 0.03 | 0.02 | 1.28 | 2.04e-01 | 7.13e-01 |
|  | REM Delta | -0.04 | 0.06 | -0.65 | 5.18e-01 | 8.06e-01 |
|  | REM Sigma | 0.04 | 0.04 | 0.83 | 4.09e-01 | 7.15e-01 |
|  | REM Theta | -0.01 | 0.05 | -0.13 | 8.93e-01 | 9.32e-01 |
|  | REM Alpha | 0.01 | 0.05 | 0.20 | 8.45e-01 | 8.45e-01 |
|  | REM Beta | 0.00 | 0.01 | 0.65 | 5.18e-01 | 6.15e-01 |
| Sleep spindle features | Integrated slow spindle activity | -0.15 | 0.12 | -1.29 | 2.03e-01 | 3.06e-01 |
|  | Integrated fast spindle activity | -0.03 | 0.09 | -0.35 | 7.25e-01 | 7.66e-01 |
|  | Slow spindle chirp | **0.25** | **0.09** | **2.82** | **6.07e-03** | **3.85e-02** |
|  | Fast spindle chirp | 0.15 | 0.10 | 1.50 | 1.39e-01 | 3.04e-01 |
|  | Slow spindle frequency | 0.16 | 0.12 | 1.34 | 1.85e-01 | 3.06e-01 |
|  | Fast spindle frequency | 0.08 | 0.12 | 0.69 | 4.90e-01 | 6.15e-01 |

Note. Only participants with at least 3 nights of data included (N=80). Abbreviations: NREM, non-rapid eye movement sleep; REM, rapid eye movement sleep. Models account for age and sex as fixed effects, and participant nested within site as a random effect. Microarchitecture variables also included channel as fixed effect. Values in bold survive correction for multiple comparisons (false discovery rate, *p*_FDR_<0.05).

Table S12. Effect of age on intra-individual variability of macro- and microarchitecture variables for at-home recordings only.

| Outcome | | β | SE | t | *p* | *p*_FDR_ |
| --- | --- | --- | --- | --- | --- | --- |
| Sleep macro-architecture features | Total Sleep Time | 0.04 | 0.11 | 0.36 | 7.20e-01 | 8.23e-01 |
|  | Wake After Sleep Onset | -0.13 | 0.07 | -1.85 | 6.91e-02 | 1.84e-01 |
|  | N2% | -0.09 | 0.12 | -0.78 | 4.37e-01 | 6.32e-01 |
|  | N3% | 0.00 | 0.08 | -0.02 | 9.83e-01 | 9.83e-01 |
|  | REM% | 0.12 | 0.08 | 1.46 | 1.49e-01 | 2.99e-01 |
|  | REM latency | **-0.40** | **0.09** | **-4.28** | **5.81e-05** | **4.65e-04** |
|  | Sleep Efficiency | -0.17 | 0.08 | -2.02 | 4.77e-02 | 1.84e-01 |
|  | Time In Bed | 0.05 | 0.07 | 0.72 | 4.74e-01 | 1.84e-01 |
|  | Transition index from NREM to REM | -0.05 | 0.12 | -0.38 | 7.07e-01 | 9.88e-01 |
|  | Total sleep cycles | -0.15 | 0.10 | -1.57 | 1.20e-01 | 6.01e-01 |
|  | Total cycle duration | 0.00 | 0.12 | -0.03 | 9.79e-01 | 9.88e-01 |
|  | Total NREM cycle duration | 0.00 | 0.11 | 0.01 | 9.88e-01 | 9.88e-01 |
|  | Total REM cycle duration | -0.06 | 0.12 | -0.45 | 6.56e-01 | 9.88e-01 |
| Sleep spindle and coupling features | Slow spindle duration | 0.10 | 0.08 | 1.16 | 2.48e-01 | 5.16e-01 |
|  | Slow spindle amplitude | -0.02 | 0.04 | -0.48 | 6.35e-01 | 9.52e-01 |
|  | Slow spindle density | -0.11 | 0.10 | -1.11 | 2.69e-01 | 6.72e-01 |
|  | Fast spindle duration | -0.13 | 0.10 | -1.35 | 1.81e-01 | 5.44e-01 |
|  | Fast spindle amplitude | 0.02 | 0.07 | 0.31 | 7.57e-01 | 9.52e-01 |
|  | Fast spindle density | -0.02 | 0.05 | -0.37 | 7.14e-01 | 9.52e-01 |
|  | Slow oscillation-slow spindle coupling magnitude | 0.11 | 0.06 | 1.79 | 7.85e-02 | 3.92e-01 |
|  | Slow oscillation-fast spindle coupling magnitude | 0.02 | 0.09 | 0.25 | 8.05e-01 | 9.52e-01 |
| Slow oscillation features | Slow oscillation duration | 0.10 | 0.08 | 1.16 | 2.48e-01 | 5.16e-01 |
|  | Median slow oscillation peak to peak amplitude | -0.11 | 0.09 | -1.17 | 2.46e-01 | 5.16e-01 |
| Relative power during NREM and REM | NREM Delta | 0.10 | 0.10 | 1.07 | 2.88e-01 | 5.16e-01 |
|  | NREM Sigma | 0.10 | 0.08 | 1.34 | 1.84e-01 | 5.16e-01 |
|  | NREM Theta | 0.10 | 0.06 | 1.67 | 9.83e-02 | 4.43e-01 |
|  | NREM Beta | 0.03 | 0.01 | 1.98 | 5.13e-02 | 4.43e-01 |
|  | REM Delta | 0.12 | 0.14 | 0.87 | 3.88e-01 | 5.16e-01 |
|  | REM Sigma | 0.05 | 0.05 | 0.88 | 3.84e-01 | 5.16e-01 |
|  | REM Theta | 0.08 | 0.09 | 0.84 | 4.01e-01 | 5.16e-01 |
|  | REM Alpha | 0.05 | 0.04 | 1.16 | 2.49e-01 | 5.16e-01 |
|  | REM Beta | 0.03 | 0.01 | 1.98 | 5.13e-01 | 4.43e-01 |
| Absolute power during NREM and REM | **NREM Delta** | **-0.29** | **0.09** | **-3.37** | **1.24e-03** | **1.85e-02** |
|  | NREM Sigma | 0.01 | 0.04 | 0.14 | 8.88e-01 | 9.52e-01 |
|  | NREM Theta | -0.02 | 0.06 | -0.36 | 7.19e-01 | 9.52e-01 |
|  | NREM Beta | 0.00 | 0.02 | 0.02 | 9.82e-01 | 9.82e-01 |
|  | REM Delta | -0.15 | 0.08 | -1.94 | 5.67e-02 | 3.92e-01 |
|  | REM Sigma | -0.01 | 0.04 | -0.29 | 7.75e-01 | 9.52e-01 |
|  | REM Theta | -0.01 | 0.07 | -0.18 | 8.56e-01 | 9.52e-01 |
|  | REM Alpha | -0.03 | 0.05 | -0.59 | 5.60e-01 | 6.30e-01 |
|  | REM Beta | 0.00 | 0.00 | -0.94 | 3.50e-01 | 5.16e-01 |
| Sleep spindle features | Integrated slow spindle activity | -0.19 | 0.11 | -1.68 | 9.79e-01 | 4.43e-01 |
|  | Integrated fast spindle activity | -0.06 | 0.09 | -0.71 | 4.82e-01 | 5.78e-01 |
|  | Slow spindle chirp | 0.21 | 0.08 | 2.48 | 1.54e-02 | 2.77e-01 |
|  | Fast spindle chirp | 0.04 | 0.09 | 0.40 | 6.89e-01 | 6.89e-01 |
|  | Slow spindle frequency | 0.10 | 0.09 | 1.12 | 2.66e-01 | 5.16e-01 |
|  | Fast spindle frequency | 0.06 | 0.12 | 0.53 | 5.99e-01 | 6.34e-01 |

Note. Participants with at least 2 nights of data included (N=75). Abbreviations: NREM, non-rapid eye movement sleep; REM, rapid eye movement sleep. Models account for age and sex as fixed effects, and participant nested within site as a random effect. Microarchitecture variables also included channel as fixed effect. Values in bold survive correction for multiple comparisons (false discovery rate, *p*_FDR_<0.05).

Table S13. Within-subjects comparisons of macro- and microarchitecture variables between at-home and in-lab recordings.

| Outcome | | Mean Difference (Home-Lab) | t | df | *p* | *p*_FDR_ |
| --- | --- | --- | --- | --- | --- | --- |
| Sleep macro-architecture features | Total Sleep Time | 8.17 | 0.52 | 32 | 6.09e-01 | 6.09e-01 |
|  | Wake After Sleep Onset | -5.31 | -0.77 | 32 | 4.49e-01 | 6.09e-01 |
|  | N2% | 1.33 | 1.08 | 32 | 2.87e-01 | 6.09e-01 |
|  | N3% | -2.65 | -2.01 | 32 | 5.34e-02 | 4.27e-01 |
|  | REM% | 1.33 | 0.92 | 32 | 3.66e-01 | 6.09e-01 |
|  | REM latency | 5.34 | 0.65 | 32 | 5.23e-01 | 6.09e-01 |
|  | Sleep Efficiency | 1.14 | 0.53 | 32 | 6.00e-01 | 6.09e-01 |
|  | Time In Bed | -17.06 | -0.93 | 32 | 3.59e-01 | 6.09e-01 |
|  | Transition index from NREM to REM | 0.01 | 1.54 | 32 | 1.34e-01 | 6.70e-01 |
|  | Total sleep cycles | 0.03 | 0.15 | 32 | 8.85e-01 | 8.85e-01 |
|  | Total cycle duration | 7.66 | 0.46 | 32 | 6.52e-01 | 8.85e-01 |
|  | Total NREM cycle duration | 2.78 | 0.24 | 32 | 8.12e-01 | 8.85e-01 |
|  | Total REM cycle duration | 5.11 | 0.72 | 32 | 4.76e-01 | 8.85e-01 |
| Sleep spindle and coupling features | Slow spindle duration | -0.02 | -1.62 | 32 | 1.16e-01 | 8.33e-01 |
|  | Slow spindle amplitude | 2.46 | 1.01 | 32 | 3.21e-01 | 8.33e-01 |
|  | Slow spindle density | -0.03 | -0.65 | 32 | 5.23e-01 | 8.72e-01 |
|  | Fast spindle duration | -0.01 | -0.87 | 32 | 3.93e-01 | 8.33e-01 |
|  | Fast spindle amplitude | 0.12 | 0.08 | 32 | 9.39e-01 | 9.92e-01 |
|  | Fast spindle density | -0.04 | -1.21 | 32 | 2.34e-01 | 8.33e-01 |
|  | Slow oscillation-slow spindle coupling magnitude | 0.00 | 0.01 | 32 | 9.92e-01 | 8.33e-01 |
|  | Slow oscillation-fast spindle coupling magnitude | 0.01 | 0.55 | 32 | 5.84e-01 | 8.76e-01 |
| Slow oscillation features | Slow oscillation duration | -0.01 | -1.26 | 32 | 2.18e-01 | 5.91e-01 |
|  | Median slow oscillation peak to peak amplitude | -0.03 | -0.01 | 32 | 9.95e-01 | 9.95e-01 |
| Relative power during NREM and REM | NREM Delta | -0.01 | -1.52 | 32 | 1.38e-01 | 5.41e-01 |
|  | NREM Sigma | 0.00 | 0.92 | 32 | 3.63e-01 | 6.81e-01 |
|  | NREM Theta | 0.01 | 1.42 | 32 | 1.65e-01 | 5.41e-01 |
|  | NREM Beta | 0.00 | -0.29 | 32 | 7.77e-01 | 8.78e-01 |
|  | REM Delta | 0.00 | 0.27 | 32 | 7.86e-01 | 8.78e-01 |
|  | REM Sigma | -0.00 | -0.86 | 32 | 3.94e-01 | 6.81e-01 |
|  | REM Theta | 0.01 | 1.02 | 32 | 3.17e-01 | 6.70e-01 |
|  | REM Alpha | -0.00 | -0.67 | 32 | 5.09e-01 | 6.91e-01 |
|  | REM Beta | -0.00 | -1.05 | 32 | 3.01e-01 | 6.70e-01 |
| Absolute power during NREM and REM | NREM Delta | 13.53 | 0.30 | 32 | 7.69e-01 | 9.92e-01 |
|  | NREM Sigma | 0.83 | 0.80 | 32 | 4.32e-01 | 8.33e-01 |
|  | NREM Theta | 13.94 | 0.79 | 32 | 4.37e-01 | 8.33e-01 |
|  | NREM Beta | -0.09 | -0.01 | 32 | 9.48e-01 | 9.92e-01 |
|  | REM Delta | 12.57 | 0.16 | 32 | 8.77e-01 | 9.92e-01 |
|  | REM Sigma | 1.36 | 0.78 | 32 | 4.44e-01 | 8.33e-01 |
|  | REM Theta | 27.49 | 1.36 | 32 | 1.84e-01 | 8.33e-01 |
|  | REM Alpha | 4.37 | 0.73 | 32 | 4.70e-01 | 6.91e-01 |
|  | REM Beta | 0.61 | 0.16 | 32 | 8.75e-01 | 9.24e-01 |
| Sleep spindle features | Integrated slow spindle activity | -34.61 | -1.57 | 32 | 1.26e-01 | 5.41e-01 |
|  | Integrated fast spindle activity | -19.18 | -1.40 | 32 | 1.71e-01 | 5.41e-01 |
|  | Slow spindle chirp | -0.04 | -2.24 | 32 | 3.20e-02 | 5.41e-01 |
|  | Fast spindle chirp | -0.02 | -0.71 | 32 | 4.85e-01 | 6.91e-01 |
|  | Slow spindle frequency | -0.02 | -0.54 | 32 | 5.92e-01 | 7.50e-01 |
|  | Fast spindle frequency | -0.06 | -1.66 | 32 | 1.07e-01 | 5.41e-01 |

Abbreviations: NREM, non-rapid eye movement sleep; REM, rapid eye movement sleep.

Figure S1. Significant associations between sleep macro-architecture and age, separated by sex.

For visualization purposes, we averaged data points across channel and night for each individual. (A) N2% sleep, (B) N3% sleep, (C) rapid-eye movement (REM) latency, (D) time in bed. Dashed line = average age effect, red = female, blue = male. Shading represents 95% confidence intervals.

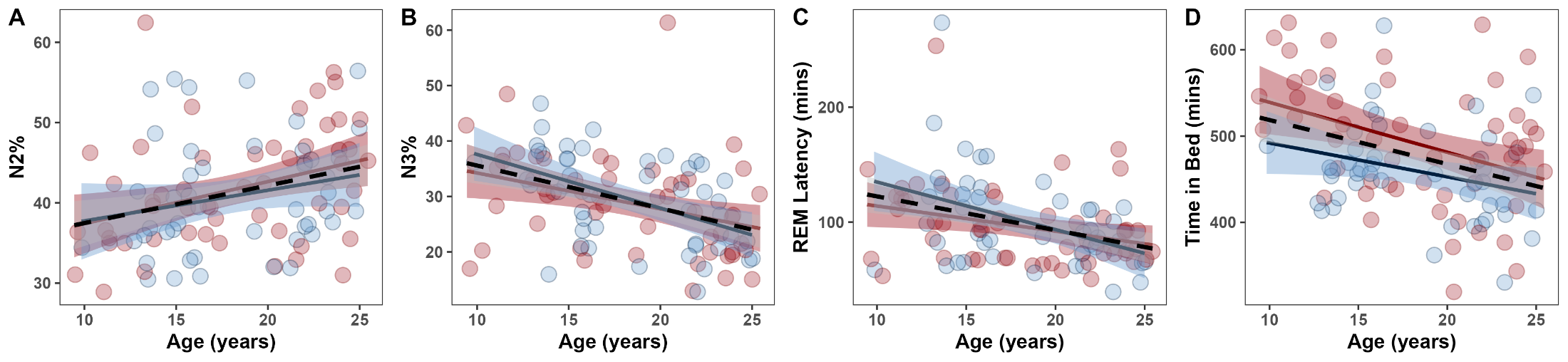

Figure S2. Associations between sleep micro-architecture and age, separated by sex.

For visualization purposes, we averaged data points across channel and night for each individual. (A) non-rapid eye movement (NREM) absolute delta power density, (B) NREM absolute theta power density, (C) slow spindle duration, (D) slow spindle density, (E) fast spindle duration, (F) slow oscillation (SO)-slow spindle coupling magnitude, (G) SO-fast spindle coupling magnitude. Dashed line = average age effect, red = female, blue = male. Shading represents 95% confidence intervals.

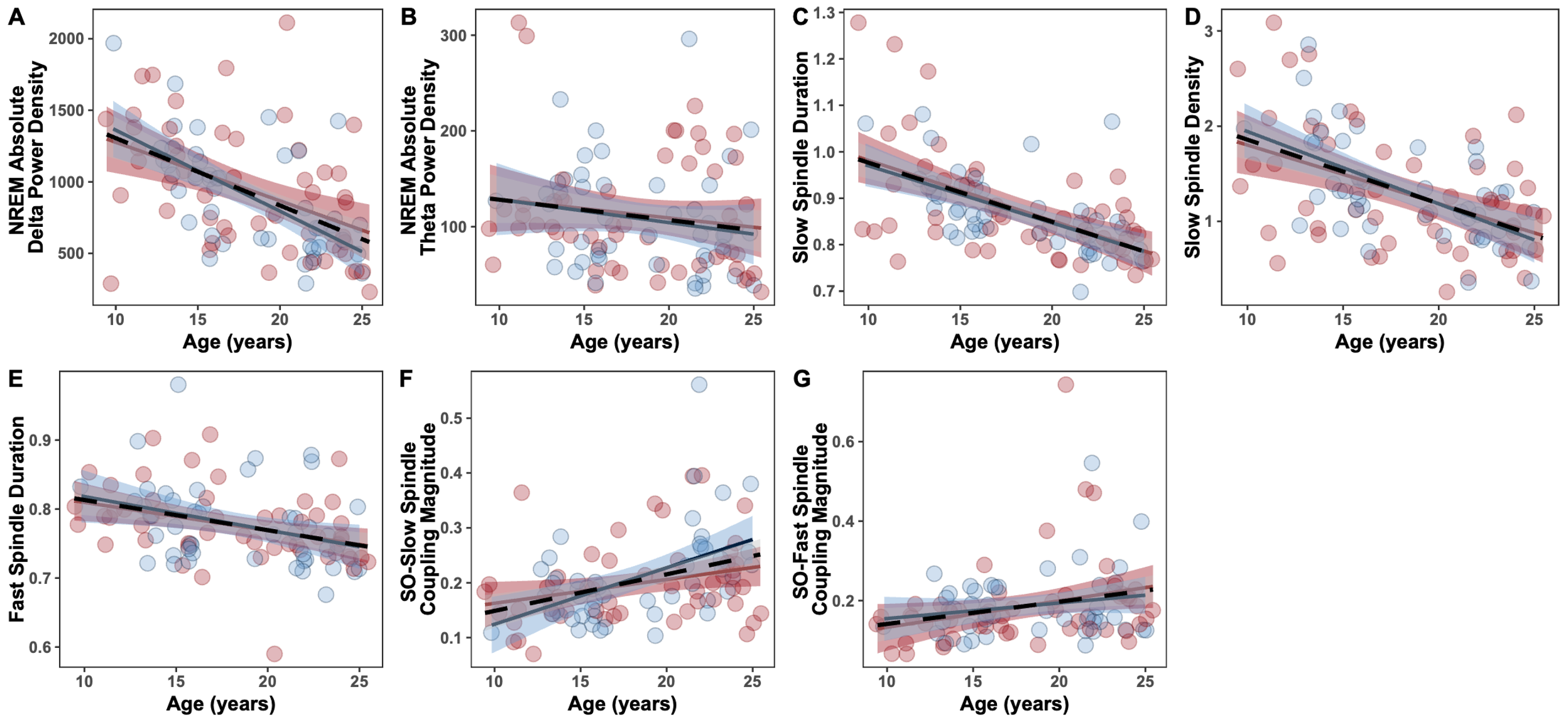

Figure S3. Macro- and micro-architecture variables showing significant sex differences.

For visualization purposes, we averaged data points across channel and night for each individual. (A) Total number of cycles, (B) REM relative sigma power density, (C) REM relative theta power density, (D) REM relative beta power density. Dashed line = average age effect, red = female, blue = male. Shading represents 95% confidence intervals. No variables show significant associations with age.

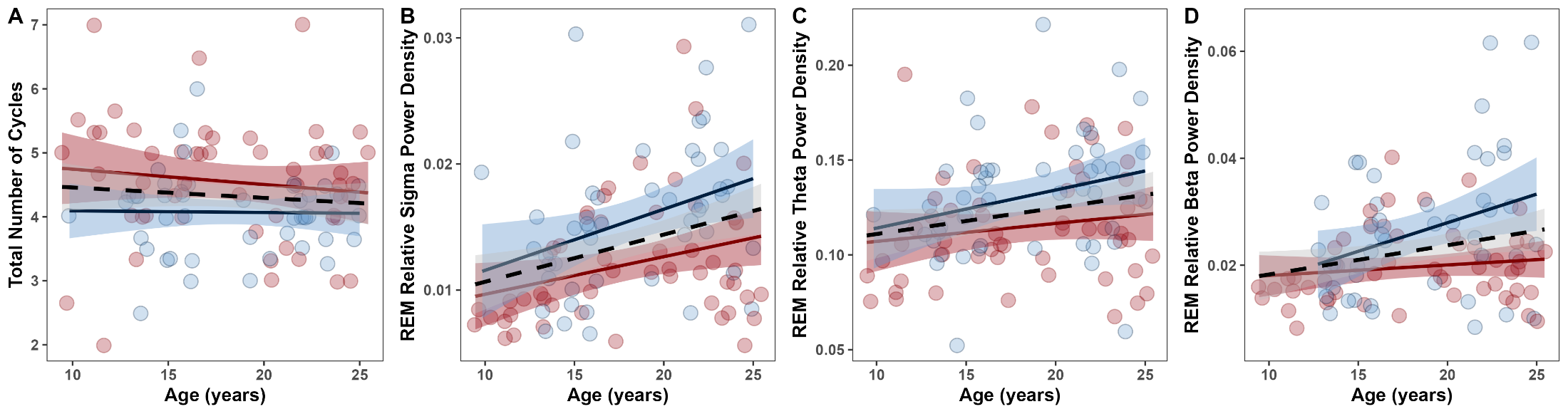

Figure S4. Age-related associations in micro-architecture variables not replicated in the present study. Data points represent average participant values across channel and night. (A) non-rapid eye movement (NREM) absolute sigma power density, (B) NREM absolute beta power density, (C) NREM absolute delta power density, (D) NREM absolute sigma power density, (E) NREM absolute theta power density, (F) slow spindle amplitude, (G) fast spindle amplitude, (H) fast spindle density. Shading represents 95% confidence intervals.

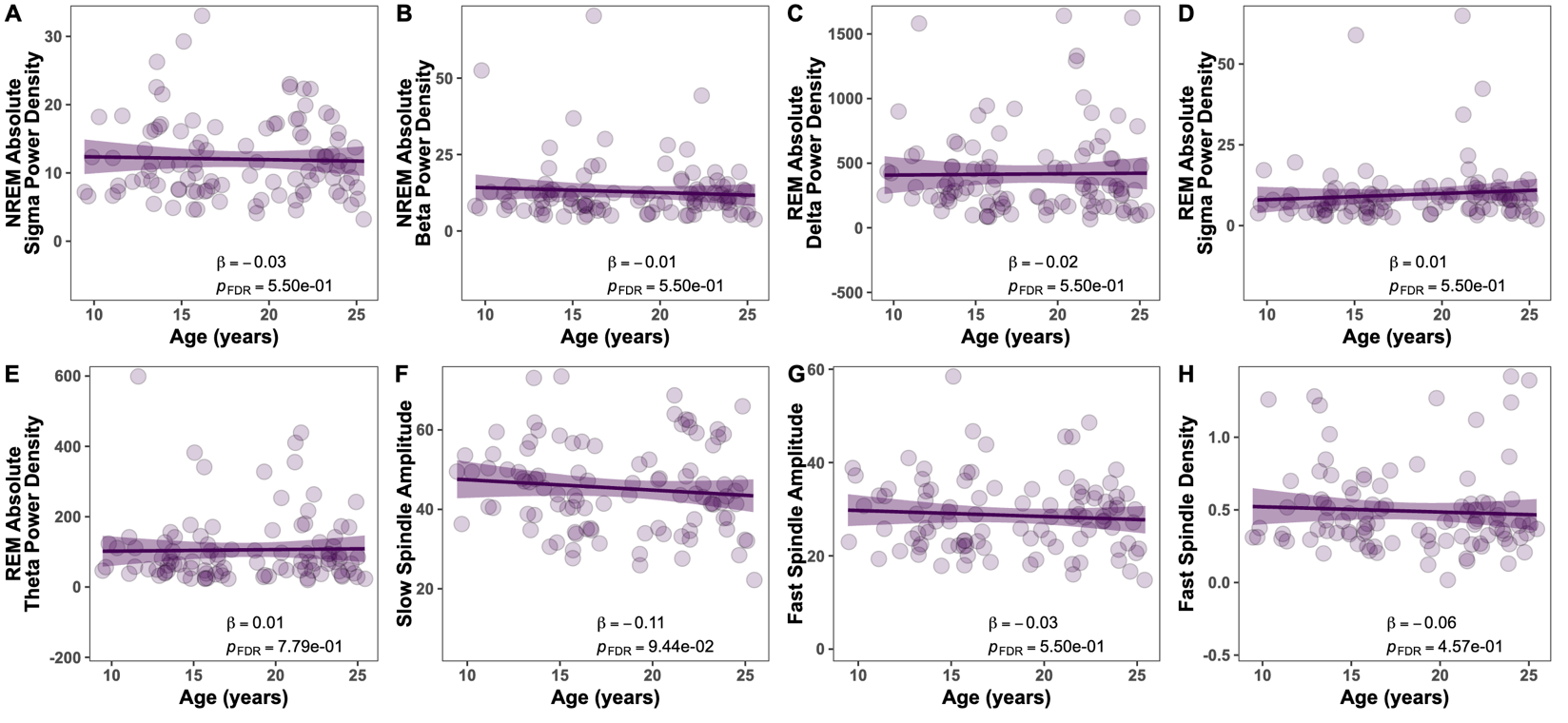
**Figure S5.** **Significant associations between age and intra-individual variability of macro- and micro-architecture variables.**

Data points represent average participant values across channel and night. (A) Intra-individual variability (IIV) in REM%, (B) IIV in SO-slow spindle coupling magnitude, (C) IIV in NREM relative theta power density, (D) IIV in NREM relative beta power density, (E) IIV in slow spindle chirp. Shading represents 95% confidence intervals.

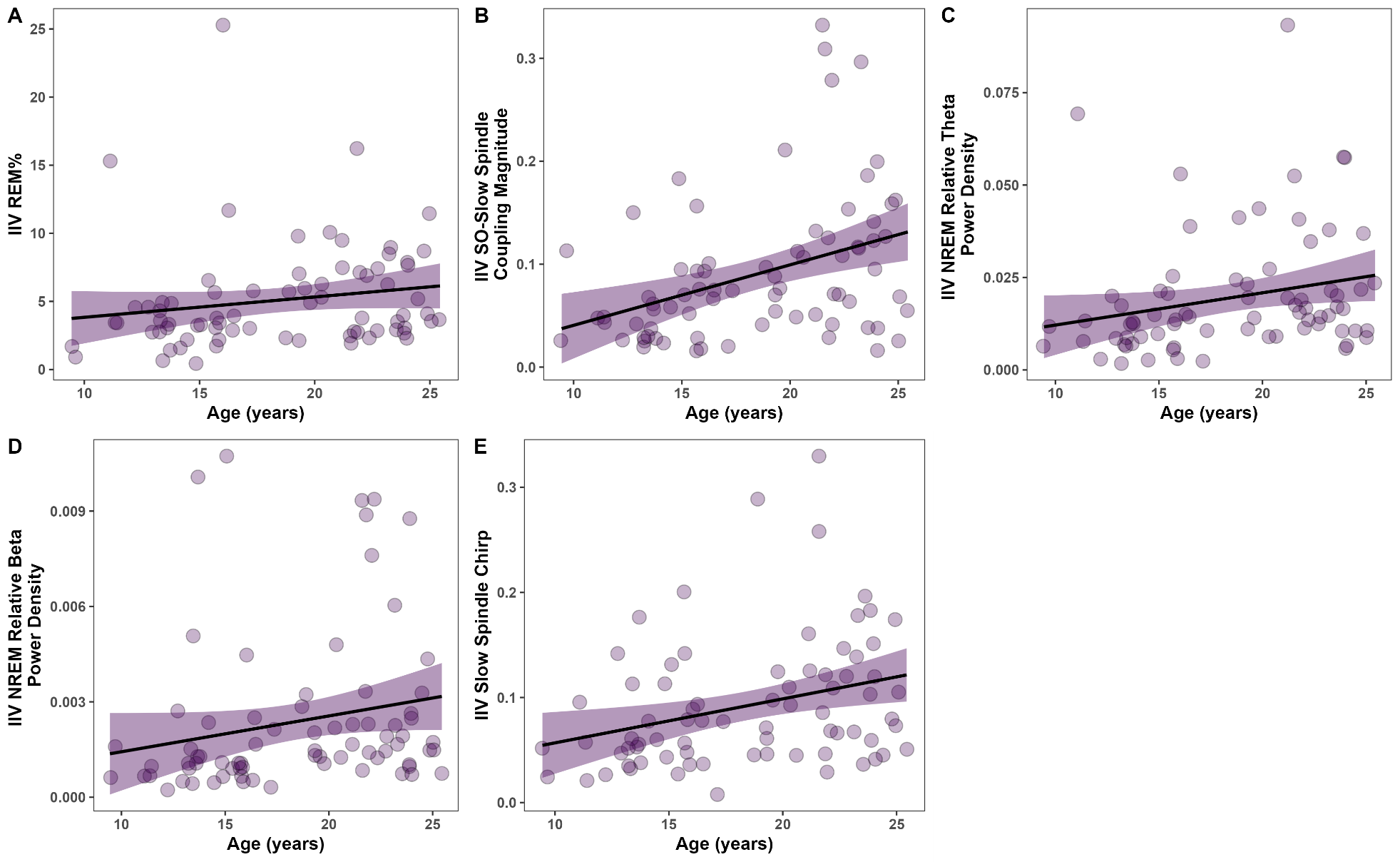

Supplemental References.
